# Mesoscale Soft Robotics for Mechanically Perturbing and Functionally Monitoring Responses in Human Organoids

**DOI:** 10.64898/2026.10.28.762586

**Authors:** Woo-Youl Maeng, Zengyao Lv, Yasaman Samei, Kyoung-Ho Ha, Kyung Su Kim, Lakshmi P. Prabakar, Ivana A. Chychula, Vasant Iyer, Binjie Li, Yuxuan Mao, Shengwei Xu, Tae Wan Park, Shupeng Li, Cory W. Holgren, Young Jin Jo, Hak-Young Ahn, Naijia Liu, Seonggwang Yoo, Seyong Oh, Jahyun Koo, Elizabeth M. McNally, Igor R. Efimov, Dominic E. Fullenkamp, Yonggang Huang, John D. Finan, John A. Rogers

## Abstract

Human organoids represent promising models for studies of significance to the biology and medical communities. Of the existing approaches for monitoring organoids, none offers the ability to apply programmable mechanical forces while simultaneously recording multi-modal physiological responses, of relevance to phenomena spanning from traumatic brain injuries to cardiac rhythm disorders. Here, we present a mesoscale soft robotic organoid interface that enables programmable delivery of forces to organoids of diverse shapes, and to map electrophysiological activity and mechanical responses. The platform combines actuation elements with integrated sensors for quantitative measurement and control. Studies using this technology reveal that spheroidal cortical organoids exhibit distinct responses to mechanical trauma, including silencing, recovery, and changes in neuronal firing patterns. Additional experiments on cardiac ring tissues demonstrate that sustained stretching induces beat-to-beat rhythm variability and altered conduction. The results establish a framework for investigating force-triggered transitions between physiological and disease-like states in human tissue models.

**HIGHLIGHTS:**

◦ A mesoscale soft robotic interface delivers programmable forces to human organoids
◦ Versatile geometries deliver compression, stretch, and shear to varied tissue shapes
◦ Real-time recording captures trauma-induced silencing and recovery in organoids
◦ Multimodal readout tracks rhythmic changes in engineered heart tissue under loading

## INTRODUCTION

Mechanical forces are critical determinants of tissue function and dysfunction, particularly in mechanically sensitive organs such as the brain ^1,2^, heart ^3^, and gastrointestinal system ^4^, where acute or chronic perturbations can trigger long-lasting pathological states. Traumatic brain injury, cardiac rhythm disorders, and other mechanically induced conditions arise from dynamic interactions between physical forces and cellular networks ^5,6^, reflecting fundamental mechanobiological principles underlying disease. Direct investigations of these processes in human tissues are, however, highly constrained due to safety and ethical limitations, thereby restricting mechanistic understanding and therapeutic development to animal models of only indirect relevance to patient care ^2,7^. This limitation has motivated the development of new approach methodologies (NAMs), human-derived experimental systems that serve as alternatives to animal models for investigating physiology and disease ^8^. By using human cells and tissues directly, NAMs capture features of human biology that animal systems cannot fully reproduce, including patient-specific genetic backgrounds, human-specific cellular responses, and developmental processes unique to human tissues.

Human organoids derived from human pluripotent stem cells provide a powerful platform for modeling tissue-level physiology *in vitro* ^9,10^. These systems reproduce key structural and functional features of native tissue, including coordinated electrical activity in cortical organoids ^11,12^ and rhythmic contractions in cardiac muscle rings ^13–15^. Optical imaging methods and recently introduced three-dimensional (3D) multielectrode arrays (MEAs) serve as the primary means for experimental investigation of real-time, functional behaviors of these systems ^16–18^. These and other existing approaches lack, however, any ability to impose controlled mechanical perturbations to the tissues while simultaneously measuring their responses. Current methods for mechanical stimulation, including bulk compression ^19^, substrate deformation ^20^, and microscale indentation ^21,22^, provide limited control over three-dimensional force application and are not readily compatible with continuous electrophysiological measurements with 3D or two-dimensional (2D) MEAs. As a result of these engineering limitations, studies of transitions from physiological to disease-like states under mechanical loading are difficult or impossible to perform on millimeter-scale human tissue models.

Here, we present a mesoscale soft robotic organoid interface (SROI) that enables programmable mechanical perturbation of 3D organoids of various shapes and sizes, while simultaneously measuring functional responses. Millimeter and sub-millimeter scale collections of actuators, acting independently or in a coordinated fashion, deliver user-defined levels of mechanical compression to spheroidal brain organoids and radially oriented tensile forces to cardiac ring tissues, all under closed-loop control. Throughout these mechanical manipulations, these systems can capture electrophysiological signals and strain distributions associated with mechanically driven transitions from physiological to disease-like states **(Figure 1A)**, continuously and in real-time.

**Figure 1.**
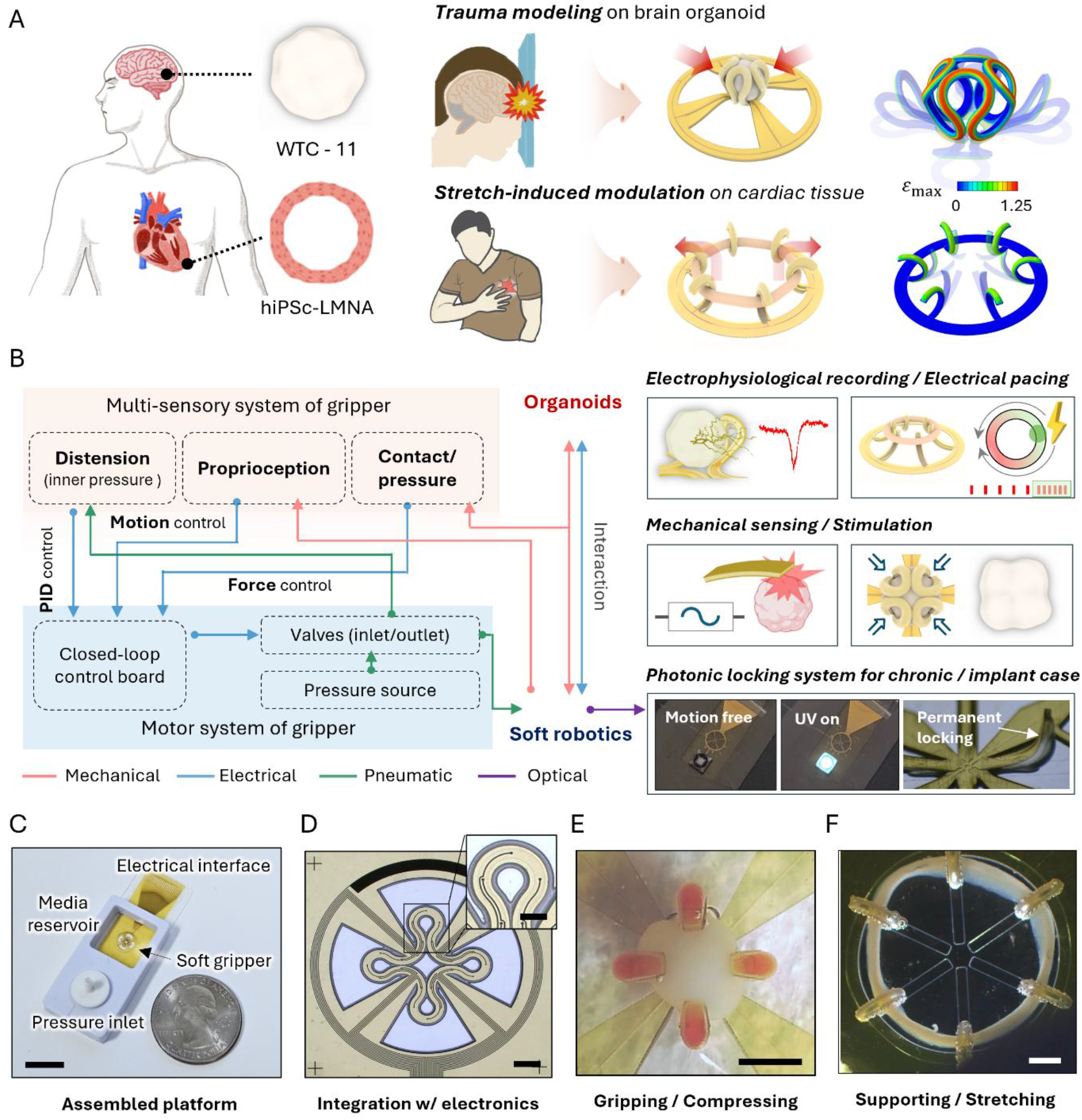
Instrumented, millimeter-scale soft robotic interfaces for modeling mechanical disease states in human organoids. **(A)** Mechanical disease modeling in two types of human organoids. Spheroidal cortical organoids (WTC-11) for traumatic brain injury and ring-shaped engineered heart tissue (hiPSc-LMNA) for mechanical perturbation are subjected to mechanical loading using instrumented, millimeter-scale soft robotic interfaces. Finite element analysis (FEA) illustrates the actuation mechanics of each system, including a device designed to apply transient compressive mechanical loading to brain organoids and one to deliver controlled stretching to cardiac organoids. **(B)** System-level schematic illustration of the multisensory soft robotic platform. The diagram depicts integrated mechanical, electrical, pneumatic, and optical pathways for interactions with organoids, via both sensing and actuation. **(C)** Image of a compact, miniaturized integrated platform (scale bar, 5 mm). **(D)** Top view of the multilayer thin-film electronics and soft actuator with aligned layouts (scale bar, 1 mm; inset, 0.5 mm). Soft robotic interface **(E)** gripping and compressing a spheroidal brain organoid (scale bar, 1 mm) and **(F)** supporting and stretching a ring-shaped cardiac organoid (scale bar, 10 mm).

In this work, we report studies enabled by this SROI platform revealing that cortical organoids exhibit distinct responses to mechanical trauma, including transient signal silencing, recovery, and cumulative alterations in neural network dynamics under repeated loading. In cardiac tissues, controlled stretching disrupts rhythmic activity and alters electrophysiological dynamics, providing a basis for modeling mechanically induced arrhythmia-like behavior. The results define a broad set of experimental capabilities for evaluating force-mediated tissue states that are inaccessible to existing approaches.

## RESULTS

### Soft robotic organoid interface (SROI)

The SROI integrates a conformal, mechanically active structure with embedded electronics as the basis for simultaneous physical interactions and electrophysiological interfaces and applied it to organoids with diverse shapes with dimensions in the millimeter and sub-millimeter regimes **(Figure 1B).** In experiments reported here, pneumatically actuated elements, operating independently or in coordination under precise control, to enable programmable delivery of compression to spheroidal brain organoids and radially oriented stretching to cardiac ring tissues. With this system, real-time monitoring of electrical activity and mechanical strains provides direct readout of functional behaviors during force delivery. Finite element analysis (FEA) informs the design of these structures and enables quantitative interpretation of interactions with the human organoids.

The SROI consists of a soft elastomeric structure that supports a network of sealed channels (silicone; 80µm channel layer, and 30 µm base layer) and an integrated system of thin, flexible electronics (5 µm polyimide substrate/10 nm Cr /200 nm Au/ 5 µm polyimide encapsulation) **(Figure S1A)**. Hydraulic and electrical connections to a proportional-integral-derivative (PID) controlled pneumatic valve system **(Figure S2)** with on-board electronics at the perimeter regions of the device. Upon pumping of air into the channels, the elastomeric structures stretch such that a modulus mismatch between the layered structure of the system drives non-linear deformations along predefined trajectories, as the basis for controlled, reversible actuation^23^. The electronics enable electrophysiological signal recording, electrical pacing, proprioceptive sensing of soft robotic deformation, and real-time monitoring of contact states at the organoid-device interface. These collective inputs enable closed-loop feedback approaches that couple mechanical, electrical, and pneumatic interactions, for real-time autonomous regulation of the injected pneumatic volume. This feedback loop maintains stable, dynamic interactions between the organoid and the soft robotic interface, in a manner defined by the user. Furthermore, for chronic organoid monitoring or implantable tissue applications, photocurable liquid precursors can be injected into the pneumatic channels and then cured into solid form to lock the device geometry into a desired state, thereby allowing operation in either an active actuation mode or a structurally fixed mode, depending on the application.

The platform includes a pneumatic inlet for pressure delivery and an electronic interconnect for electrical interfacing, and a surrounding layer of poly(dimethyl siloxane) that forms a reservoir for the organoid and culture medium **(Figure 1C)**. The fabrication process relies on precise alignment of the soft robotic layer with the flexible electronics layer **(Figure S1B),** which in the examples presented here includes 20 electrodes (40 µm diameter) for electrophysiological recording **(Figure 1D)**. The device morphology can be tailored over a wide range, to realize actuatable cavity-type structures or articulated supports for organoids with diverse shapes and with sizes in the single millimeter range or larger. Representative images show a linear four-arm structure designed to cradle a cortical organoid with a diameter of 1.5 mm **(Figure 1E, Video 1)**, and a configuration that supports a cardiac ring tissue with inner diameter of 5.5 mm **(Figure 1F)**.

### Design versatility and motion characterization

The miniaturized soft arms within these structures (widths 300 µm) undergo large, non-linear bending deformations upon pneumatic actuation, reaching maximum strains of ∼1.19 (i.e., 119%) in the elastomeric component of the system during maximum extension **(Figure S3C)**. FEA simulations employing an incompressible, second-order Yeoh hyperelastic model ^24^, with material parameters obtained from uniaxial tests **(Figure S3D)**, and this system accurately capture this behavior and guide geometric designs tailored to targeted organoid geometries and tissue morphologies. Coordinated actuation of multiple arms defines a closed, inward-curving structure that forms a near-spherical enclosure. The extent of geometric conformity depends on the number and arrangement of the arms, with increased arm counts yielding more uniform and continuous contact profiles, as supported by FEA simulations **(Figure 2A, Figure S4)**. This configuration enables controlled and reproducible mechanical coupling with organoids of varying shapes and sizes (**Video 2**). Different layouts, including linear design (four-arm), petal-shaped design (six-arm), and full-cover (spider-like) geometries modulate the arm-organoid contact coverage from 43% to 100%. Beyond spheroidal geometries, these designs extend to tissues with diverse morphologies, including dual cavity architectures for tissue-tissue interfaces in assembloids, linear structures that could accommodate intestinal tissues and ring-shaped constructs suitable for cardiac or muscle rings enable delivery of compressive, tensile, and shear-compressive mechanical stimuli in programmable patterns. In addition, multi-array configurations allow parallel actuation of multiple organoids at once using a single pneumatic input, highlighting the scalability of the platform. These representative design examples demonstrate the broad range of soft robotic interfaces that can be achieved, depending on required mechanical stimulation modes (compression, shear, and stretching) and organoid shapes, dimensions, and pathology-relevant loading conditions.

**Figure 2.**
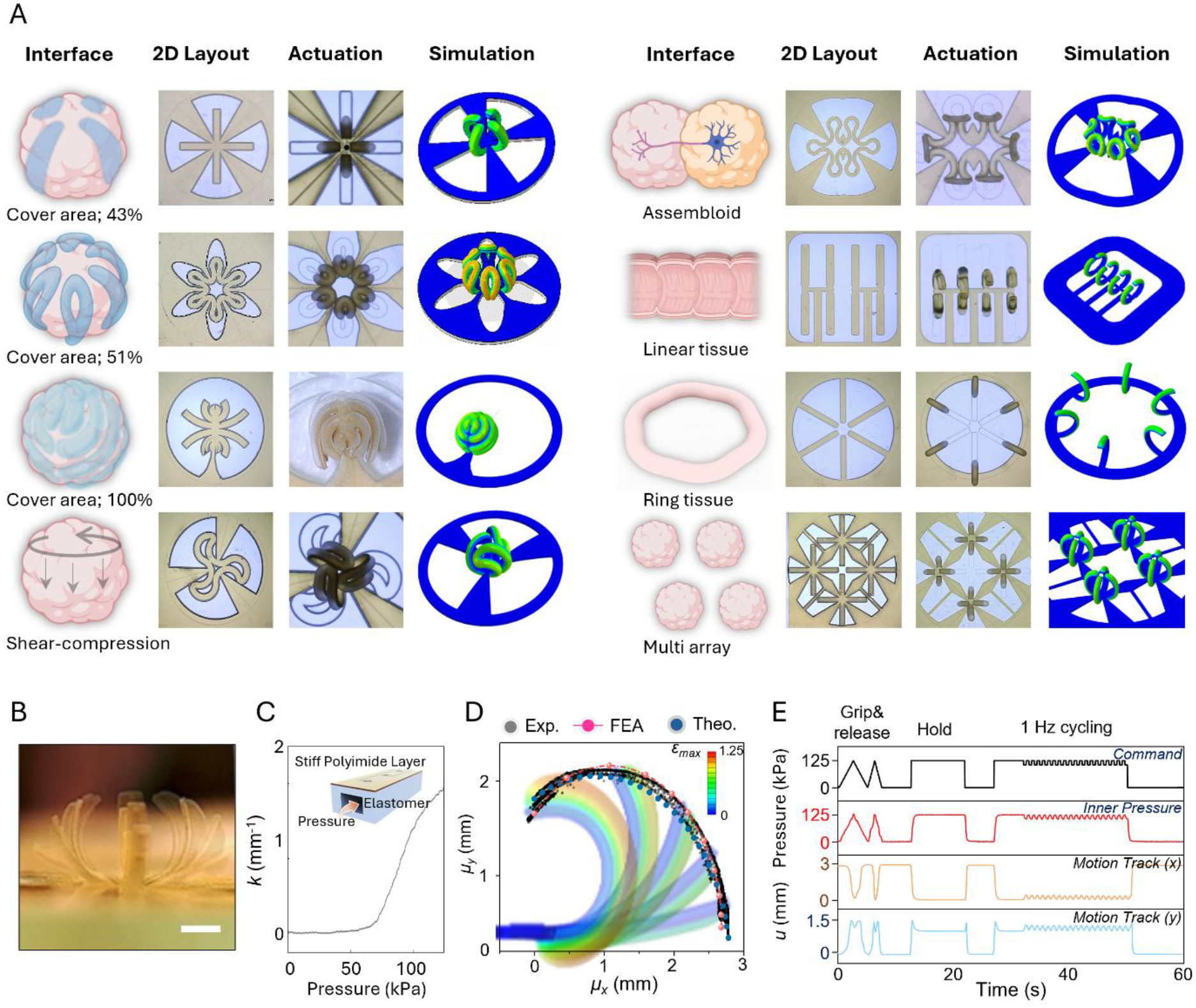
Design diversity and characteristics of instrumented, millimeter-scale soft robotic systems as organoid interfaces. **(A)** Design concepts, three-dimensional structures, and modeling of soft robotic interfaces tailored to diverse organoid pathologies, including designs for gripping spheroidal organoids, applying shear-compressive forces to them, constructing and manipulating assembloids, examining organoids in linear and ring geometries, and engaging in multi-array configurations. **(B)** Side view of the pressure-dependent actuation of a system with a four-linear arm design (Scale bar, 1mm). **(C)** Cross-sectional schematic illustration of the multilayer soft robotic structure and corresponding dependence of the curvature of a robotic arm on the inner pressure. **(D)** Side-view comparison of robotic arm tip trajectories obtained from video analysis (black), finite element analysis (FEA) (red), and circular-arc geometric calculations (blue). **(E)** Correlation between a sequence of operations applied through a micro PID valve, the measured inner pressure response in the robotic channel, and the results of side-view tracking of the gripper tip under multiple actuation modes, including gripping-and-releasing, holding, and 1 Hz cycling.

Side-view profiles of the actuation process focus on an SROI with four straight arms that each continuously bends under pneumatic pressure, forming a closed, inward-curving enclosure for capture of a spheroidal organoid **(Figure 2B)**. As the pressure increases, the arms progressively close along a reproducible trajectory. **Figure 2C** presents a cross-sectional schematic illustration of the multilayer soft robotic structure with a quantitative analysis of pressure-dependent curvature of an arm. Bending initiates at ∼60 kPa, followed by a rapid increase in curvature up to 125 kPa. This pressure-curvature relationship provides a quantitative basis for predicting arm deformations and for selecting actuation pressure ranges to achieve target geometries. **Figure 2D** compares the end-tip trajectories of a robotic arm obtained from video-based motion tracking (black dot; Tracker, V6.2.0) with predictions from FEA (red dot) and analytical modeling (blue dot, **described in the Method**). Measurements span five cycles of actuation over the full range of motion, from a full spherical cavity to a flattened configuration. These three sets of results show strong agreement and repeatable operation across cycles. Side-view video recordings yield the x-y coordinates of the arm tip, confirming that the measured trajectories closely follow the commanded motion patterns **(Figure 2E)**. The platform maintains stable pressure regulation and reproducible motion during continuous cycling. Stable holding at full actuation is also possible, with fast dynamic responses under repetitive actuation at 1 Hz. Independent addressing of each of the four robotic arms (arms 1-4) is also possible, where a separate pneumatic command drives the corresponding channel pressure to a prescribed setpoint with fast settling and stable steady-state behavior. This setpoint fidelity maintains not only during single-arm actuation but also during selective and simultaneous multi-arm actuation (arms 1 and 3, 2 and 4, 1-4), without measurable pressure interference across the channels **(Figure S5)**. These fast actuation responses enable precise and temporally controlled modulation of mechanical inputs to organoids and thus quantitative evaluation of the resulting biological responses. This control also supports traumatic loading protocols on the 1 s timescale, as described subsequently in the context of cortical organoids.

### Electronic functions and control

The polyimide (PI) layer attached on the upper surface of the elastomer serves not only as a structural component that enables non-linear bending motions of the soft robotic arm, but it also serves as a substrate for (i) electrode arrays, (ii) proprioceptive sensing elements, and (iii) contact-state sensing electronics (**Figure 3A)**. One example integrates electrodes (five circular electrodes per arm; 40 µm diameter) for electrophysiology. Platinum-electroplated electrodes on traces of gold exhibit impedances below 10 kΩ at 1 kHz, for low-noise, high fidelity recording at the organoid-device interface **(Figure 3B)**. Serpentine gold traces integrated on each arm function as resistive proprioceptive sensors, in which arm bending transduces into a change in electrical resistance. In the low-pressure range, the pneumatic channels primarily inflate. Above ∼60 kPa, the deformation shifts to a bending-dominated mode that persists for pressures up to 125 kPa **(Figure 3C)**. The resistance of these serpentine traces therefore provides an electrical readout of the mechanical state of the arm, enabling continuous electrical measurements of actuation without the need for optical imaging. FEA predicts that deformation of the arm induces strain in the serpentine gold trace (3.5 µm width) that reaches ∼0.2% at 125 kPa. This deformation thus leads to experimentally observed increases in resistance under these actuation conditions.

**Figure 3.**
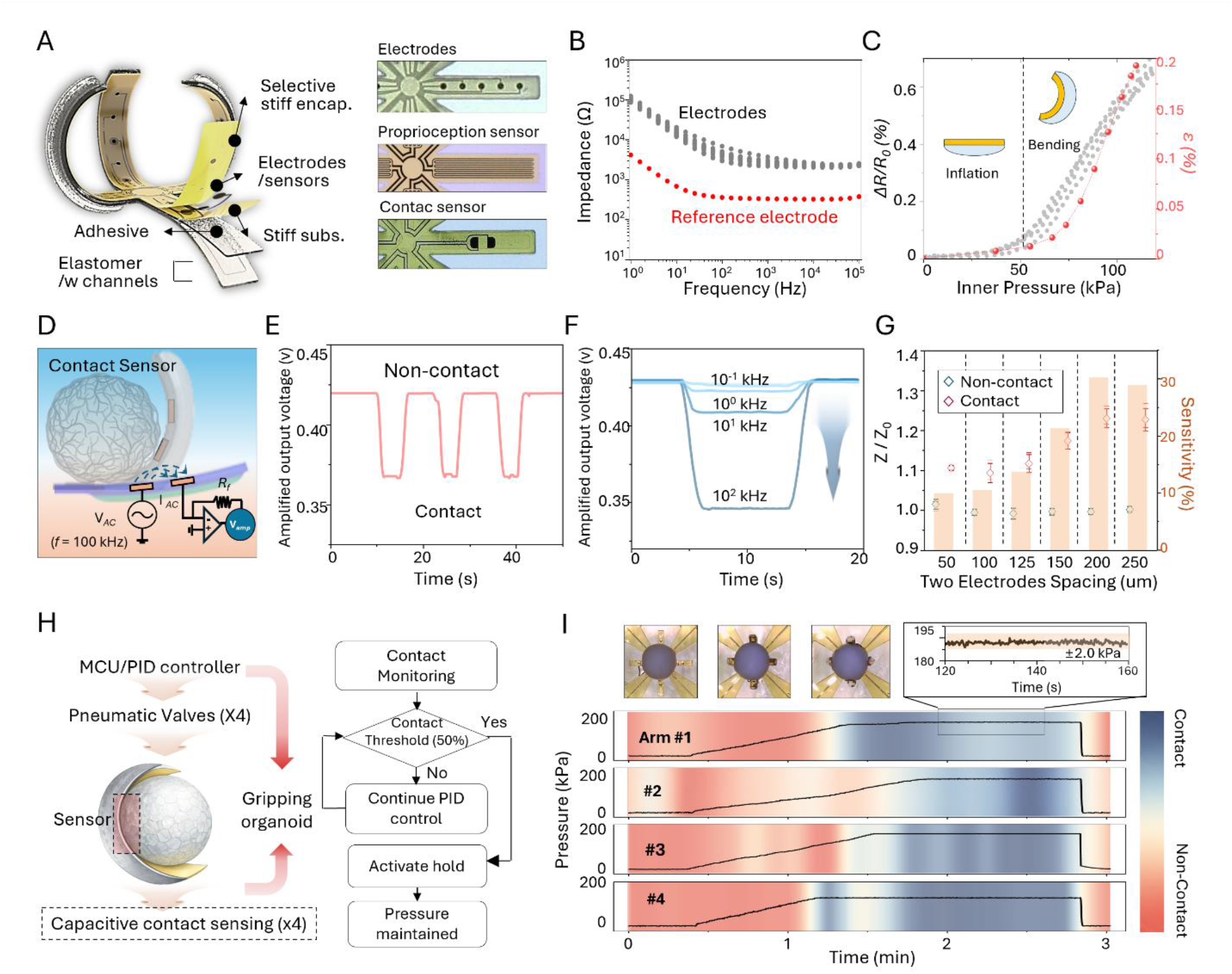
SROI platform with integrated multifunctional sensing and actuation. **(A)** Multilayer architecture of a soft gripper that integrates microfluidic actuation channels encapsulated in elastomer, thin-film electronics (PI/Cr/Au/PI), and interfacial adhesive layers. The embedded electronic components include microelectrode arrays for electrophysiological recording, a proprioceptive sensor for curvature monitoring, and contact sensors for interface detection. **(B)** Frequency-sweep impedance curves of Au electrodes coated with Pt by electrodeposition. **(C)** Normalized impedance change (Δ *R/R_0_*) of the proprioceptive sensor and corresponding simulated strain (*ε*) as a function of inner pressure. **(D)** Circuit schematic of the contact-detection sensor at the organoid-robotic arm interface. **(E)** Representative amplified output voltage response of the contact sensor under non-contact and contact conditions. **(F)** Frequency-dependent response of the contact sensor driving circuit. **(G)** Impedance responses between two electrodes under non-contact and contact conditions, along with a bar graph of corresponding sensitivity. **(H)** Overview of the closed-loop control system for conformal organoid gripping using four independently actuated robotic arms, and the corresponding control-logic flow for each arm. **(I)** Closed-loop gripping and holding operation of four independently actuated arms. Channel pressure increases until the contact-sensor output reaches a predefined threshold, defined as 50% of the voltage difference between contact and non-contact states. The pressure ramp then stops, and the corresponding channel pressure serves as the holding setpoint. Pre- and post-operation photographs confirm stable conformal contact at multiple positions around the spheroidal organoid, with pressure maintained within ±2 kPa.

Each robotic arm also incorporates exposed electrodes that function as contact sensors, enabling detection of interactions between the robotic arm and the organoids (Fig. 3D). Contact-dependent changes in impedance converted into voltage signals allow for reliable identification of contact and non-contact states **(Figure 3E)**. The sensors operate using high-frequency electrical excitation, integrated with electrophysiological measurements without significant interference. Changes in contact state produce distinct and repeatable responses during actuation, enabling dynamic tracking of arm-organoid interactions. The nature of the response depends on operating frequency and electrode spacing **(Figure 3F-G, Figure S6)**, as the basis for real-time feedback on contact conditions during actuation.

A representative SROI system includes four independently actuated robotic arms with manipulations guided by contact sensing and a microcontroller unit (MCU)-based PID control system. **Figure 3H** summarizes the overall closed-loop architecture, in which contact-sensor outputs from each arm pass to the MCU to guide control the corresponding pneumatic valves **(Figure S7)**. The control logic operates independently for each arm, allowing the system to identify contact at different positions around a spheroidal organoid and to regulate the pressure in each channel accordingly. This approach enables conformal gripping of organoids that are only approximately spherical under predefined actuation sequences. **Figure 3I** visualizes the responsiveness and holding stability of this closed-loop operation. Here, pressure gradually increases in each channel during continuous monitoring through the output of the contact-sensor. For each sensor, the contact threshold corresponds to 50% of the voltage difference between the non-contact and fully contacted states. When the sensor output exceeds this threshold, feedback from the contact sensor terminates the pressure ramp. The pressure at this moment corresponds to the holding setpoint, maintained by the PID controller through inlet and exhaust valve operation. In this state, the robotic arms remain fixed in conformal contact with the organoid, with pressure maintained within approximately ±2 kPa, as shown in the inset of **Figure 3I**. In the following organoid studies, the SROI serves as a mechanically actuated electrophysiology interface, with integrated electrodes contacting the organoid surface to record functional signals during controlled mechanical loading.

### Real-time interrogation of mechanophysiological responses in brain organoids

A first example of the unique utility of the SROI platform involves controlled delivery of traumatic mechanical loading onto cortical organoids, while continuously recording electrophysiology. Day-253 cortical organoids derived from the WTC-11 induced pluripotent stem cell (iPSC) line serve as a cortical model **(Figure 4A).** Confocal Ca^2+^ imaging further captures spontaneous network activity **(Video 3)**. The organoids (1.6-1.8 mm) exhibit high cellular density with MAP2-positive neurites and S100B-positive glial populations, confirming terminal differentiation of neurons and astrocytes, respectively. A stage top incubator mounted on the confocal microscope maintains 37℃ and provides 10 min. stabilization to minimize transient disturbances during transfer and gripping of each organoid into a 4-arm SROI platform. The electrodes connect to recorder (Intan RHD), and the PID-valve system precisely regulates gripping and mechanical loading (**Figure S8**). The setup synchronizes acquisition of electrophysiological data (20 kHz) with confocal Ca^2+^ imaging (1 Hz).

**Figure 4.**
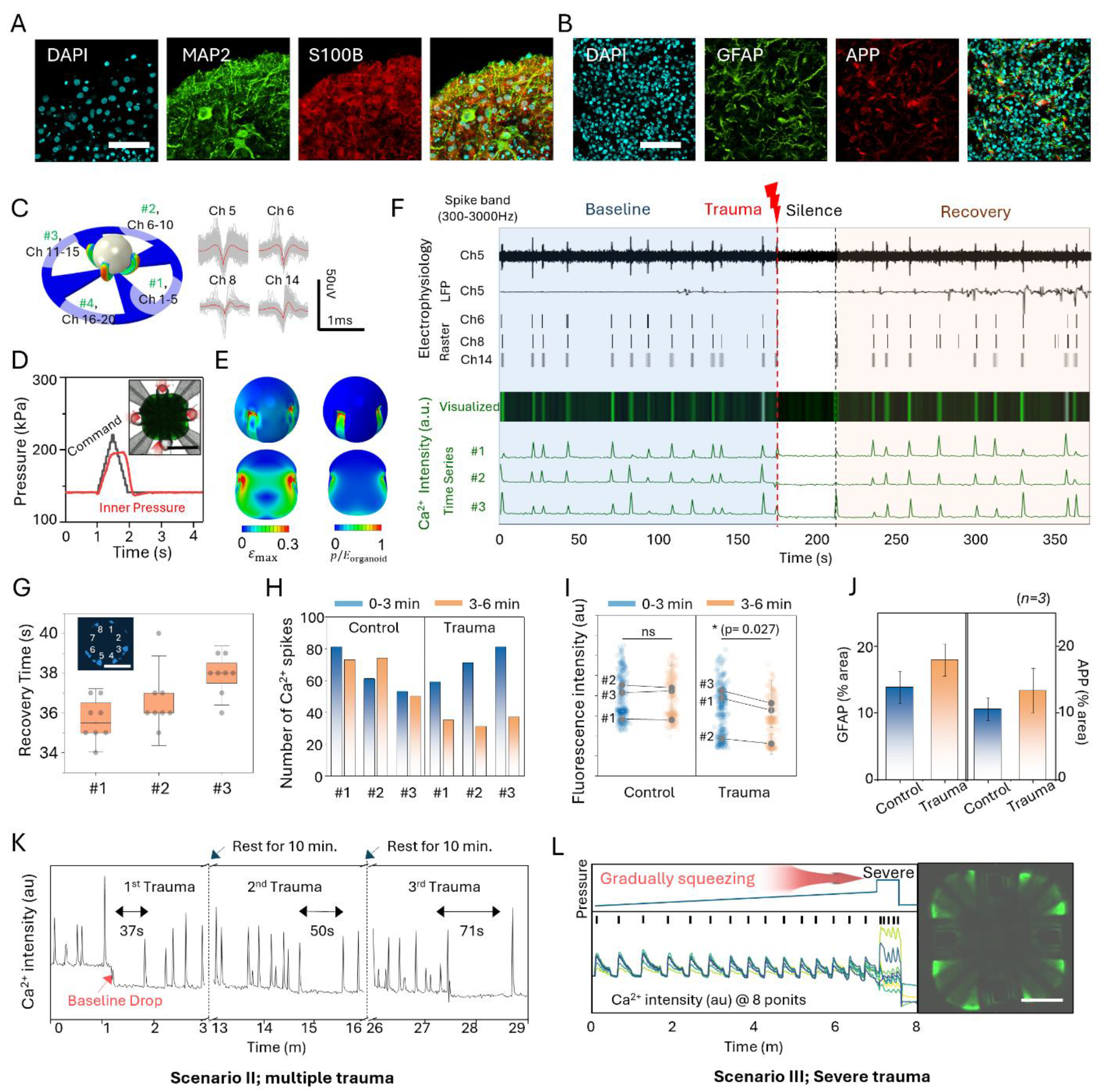
Real-time interrogation of mechanophysiology in brain organoids. Fluorescence images of WTC-11 brain organoids. **(A)** Cell-type markers; higher-magnification (20x) images stained for DAPI and MAP2 (neuronal), and S100B (glial), and merged channels (scale bar, 50μm). **(B)** Injury-associated markers; representative post-trauma fluorescence images acquired following transient mechanical impact and stained for DAPI, GFAP (astrocyte), and APP (amyloid precursor protein), and merged channels (scale bar, 50μm). **(C)** Electrode channel map and representative electrophysiology recordings from selected channels including spike waveforms from channels 5, 6, 8, and 14. **(D)** PID valve command (black) and measured pressure (red) used to deliver a 1 s transient impact (500 ms pressurization, 500 ms depressurization). Representative snapshot at impact (inset: scale bar, 1 mm). **(E)** Maximum principal strain distribution within the organoid from (top left) a side view and (bottom-left) a vertical cross section. Strain localizes at the contact area but remains strictly below 50% throughout, Pressure concentrates at the near-surface contact area, peaking at 293 Pa, which exceeds the organoid’s elastic modulus (*E_organoid_* = 200 Pa) ^27^, and suggests that the local strain is large. **(F)** Time-synchronized electrophysiological recordings and fluorescence intensity during trauma induction. Action potential (AP) and local field potential (LFP) time series from channel 5 are shown with corresponding raster plots from channels 6, 8, 14. After a 3-min baseline, a 1 s transient impact leads to silencing of both electrophysiological activity and fluorescence signals, with subsequent recovery. **(G)** Recovery time after trauma quantified across 8-predefined fluorescence regions/ROIs in three organoids (scale bar, 1 mm). **(H)** Comparison of Ca^2+^ spiking counts between control and post-trauma (*n=3*). **(I)** Comparison of fluorescence intensity pre- and post-trauma (*n=3*). **(J)** Quantification of GFAP and APP positive area (% area) from immunostained sections of fixed organoids in control and trauma groups (*n=3*). **(K)** Time series of Ca2+ responses to three multiple trauma events within a single organoid, with 10-min recovery intervals between events. **(L)** Fluorescence intensity profiles from 8-detection points during gradual compression followed by an impulsive increase to a severe compression level. (scale bar, 500 μm).

Under baseline conditions, recordings from four representative electrodes reveal spontaneous spiking with ∼20 µV amplitude and 400-500 µs duration **(Figure 4C)**. The protocol involves application of a transient traumatic load (1 s overall - 500 ms compression, 500 ms release). Confocal imaging reveals rapid deformations and geometric changes during loading **(Figure 4D)**. According to FEA, this procedure imposes localized strain at the contact area **(Figure 4E)**, with 38% of the total volume of the organoid experiencing strains of more than 10%. The strain remains strictly below 50% throughout the organoid, thus avoiding the threshold for severe traumatic axonal injury (TAI) ^25^. The experiment follows a 3-min baseline, 1-s trauma, 3-min post-trauma sequence. Figure 4D overlays synchronized electrophysiological recordings (action potentials and field potentials) with confocal Ca^2+^ fluorescence images across the experimental timeline. Electrophysiological raster plots closely coincide with Ca^2+^ signals, indicating strong concordance between these electrical and optical measurements. Immediately following the 1-s trauma, both signals exhibit transient silencing and subsequently recover in a temporally coupled manner approximately 36 s after loading **(Figure 4F, Video 4)**. This silencing-recovery pattern reproduces consistently across three trauma organoids **(Figure S9)**. Notably, local field potential (LFP; low-frequency components of extracellular potentials, 0.1-250 Hz band-pass-filtered) activity shows increased variability and bursting-like dynamics after recovery, suggesting a functional network state transition or reorganization rather than a simple return to baseline **(Figure S10)** ^11,26^. Further analysis involves quantification of Ca^2+^ fluorescence data across eight separate regions of interest (ROI) per organoid. Recovery times across these ROIs show a consistent silencing period of approximately 35-39s in all three organoids **(Figure 4G)**. Examination of Ca^2+^ spiking events over the 3-min windows before and after trauma indicates a significant reduction in spiking frequency (50±9%) relative to the behaviors of control organoids **(Figure 4H)**. The downward shift in Ca^2+^ fluorescence intensity across these ROIs indicates a global reduction in network activity following trauma, rather than localized loss of signal **(Figure 4I)**. Immunofluorescence microscopy demonstrates histological markers after trauma **(Figure 4B)**. GFAP and APP staining indicate a slight increase in marker-positive areas after trauma, the differences are not statistically significant relative to controls **(Figure 4J)**. These results are consistent with a moderate brain injury that disrupts function, but does not destroy cells.

Three transient trauma events delivered at 10 min intervals to the same organoid reveal a cumulative effect on Ca^2+^ dynamics. The first trauma reduces baseline fluorescence intensity with a recovery time of ∼37 s, whereas the second and third events progressively delay recovery to ∼50 s and ∼71 s, respectively **(Figure 4K)**. This result suggests that repeated traumatic loading prolongs recovery kinetics of the neural network. In a third demonstration, gradually increasing compressive loading produces a progressive increase in Ca^2+^ spiking frequency across all regions. Under a severe high-force grip, spiking frequency increases across all ROIs, consistent with a hyperexcitability-like state, while some regions exhibit a sharp elevation in Ca^2+^ baseline. Ca^2+^ signals are subsequently lost **(Figure 4L)**. The findings indicate that precisely controlled mechanical perturbations enable real-time monitoring of Ca²⁺ dynamics of physical trauma and associated signal transitions in neural organoids, with future potential for quantitative analysis of responses under diverse loading/unloading trajectories.

### Integrated multimodal interrogation of cardiac tissues under tensile loading

Ring-shaped engineered heart tissue (EHT) renders the complex architecture of native cardiac tissue into a tractable form for studying action potential propagation and the effects of mechanical loading ^14,27^. The circumferential propagation geometry facilitates analysis of conduction dynamics to a one dimensional problem, while the accessible inner circumference enables distributed electrode contact and uniform radial mechanical loading. The six-arm radial architecture of the SROI conforms to this geometry, enabling six distributed electrical interfaces positioned at ∼60° intervals along the inner circumference within a single device **(Figure 5A)**. The integrated six-channel electrode array records spatially distributed field potentials from spontaneous beating EHTs, enabling resolution of propagation timing across the ring structure **(Figure 5B, top)**, while also delivering biphasic electrical pacing (1 mA, 400 µs) through selected channels using the same tissue interface **(Figure 5B, bottom)**. The compact, open architecture of the SROI further allows concurrent electrical recording and optical voltage and calcium imaging on an optical mapping instrument within a single experiment **(Figure 5C, top)**. To characterize electrophysiological responses under controlled pacing, the EHT is paced at cycle length from 0.7 to 1.0s, and action potential duration at 80% repolarization (APD80) and calcium transient duration at 80% recovery (CaTD80) are quantified **(Figure 5C, bottom)**. Both APD80 and CaTD80 progressively increase at longer pacing cycle lengths, demonstrating rate-dependent electrophysiological behavior. These measurements establish a baseline for subsequent stretch-induced perturbation studies.

**Figure 5.**
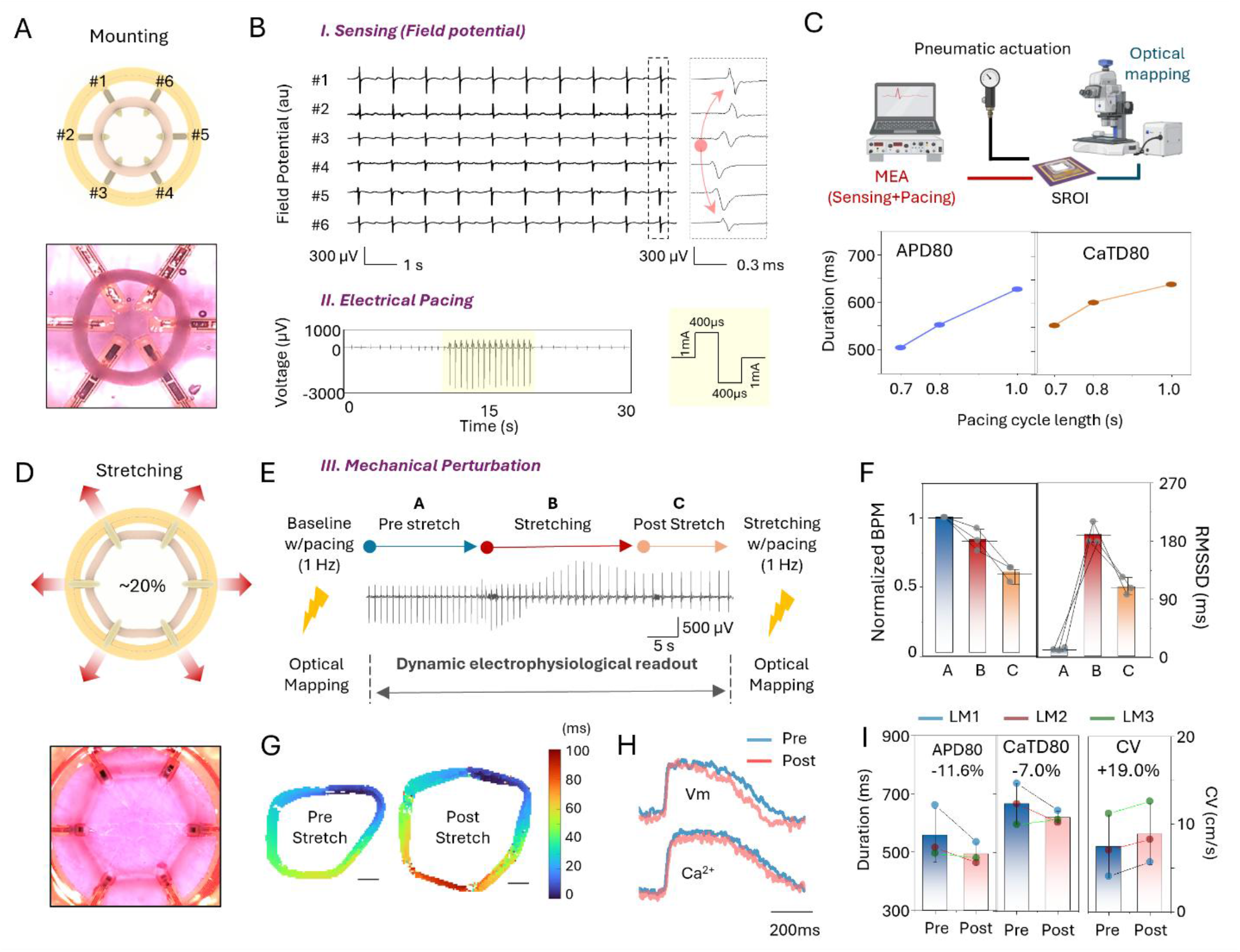
Integrated multimodal interrogation of cardiac tissues under tensile loading. **(A)** Schematic and optical image of a ring-shaped engineered heart tissue mounted on the SROI platform. The geometry enables conformal contact with six arms and spatially distributed access to the tissue. **(B)** Multi-channel field potential recordings acquired through distributed electrodes across the cardiac ring, demonstrating electrophysiological sensing and pacing within the same soft robotic interface. **(C)** Experimental configuration for concurrent electrical pacing, and optical mapping. Optical mapping quantifies rate dependent changes in APD80 and CaTD80 across pacing cycle lengths in EHTs. **(D)** Tensile loading of the ring tissue by outward pneumatic actuation of the robotic arms. Radial displacement of the arms produces circumferential stretch while maintaining tissue-electrode contact. **(E)** Dynamic field potential readout during the transition from baseline to tensile loading and post stretch. Optical mapping under electrical pacing (1Hz) is performed before and after stretching, whereas MEA recordings track real time electrophysiological changes throughout the mechanical perturbation. **(F)** Changes in normalized beat rate and beat-to-beat variability, quantified by RMSSD, across the baseline, stretching, and post-stretch phase. **(G)** Optical activation maps before and after tensile loading, showing spatial propagation patterns across the mechanically perturbed ring (scale bar: 1mm). **(H)** Representative optical voltage (Vm) and calcium (Ca^2+^) transients recorded from the EHTs before (mounted) and after (stretched) tensile loading, demonstrating stretch-associated shortening of action potential and calcium transient durations. **(I)** Optical mapping quantifies APD80, CaTD80, and conduction velocity (CV) under electrical pacing (1Hz) before and after tensile loading, revealing stretch-associated modulation of repolarization, calcium recovery, and conduction dynamics.

The SROI applies ∼20% circumferential tensile strain to the ring tissue through pneumatic actuation (path length 16.3 to 19.8 mm; +21.7%) while maintaining continuous MEA-based field potential recording throughout the stretching process **(Figure 5D, Video 5)**. Optical mapping before and after stretch enables quantitative comparison of activation and recovery dynamics. Stretch induces immediate changes in the amplitude and morphology of the field potential signal **(Figure 5E)**, demonstrating that electrophysiological dynamics under mechanical loading can be captured in real time. Quantification of spontaneous beating across pre-stretch baseline (Phase A), stretching (Phase B), and post stretch (Phase C) reveals that the beat rate decreases to 59% of baseline at Phase C, while the root mean square of successive differences (RMSSD) increases from ∼11 ms at baseline to ∼190 ms during Phase B and remains elevated at ∼109 ms during Phase C **(Figure 5F)**. These results indicate that mechanical loading produces immediate and sustained effects on spontaneous rhythm stability. Optical activation mapping under 1 Hz pacing before and after stretch capture stretch-associated changes in spatial propagation patterns across the ring tissue **(Figure 5G)**. Representative optical voltage and calcium transients further demonstrate changes in action potential and calcium waveform morphology following mechanical loading **(Figure 5H)**. Quantification from these optical recordings shows that conduction velocity (CV) increases by 19% following stretch, while APD80 and CaTD80 decrease by 11.6% and 7.0%, respectively, demonstrating that the SROI platform captures stretch-induced modulation of conduction, repolarization, and calcium recovery dynamics across tissues **(Figure 5I)**.

Electromechanical coupling is critical for normal physiology, but also where abnormal responses stretch can lead to pathology and life-threatening arrhythmias^3^. The SROI enables coupling of mechanical stretch, field potential sensing, local pacing, and optical mapping in a single platform at the tissue level. This provides a clear approach to delineating genotype and disease-specific the effects mechanical perturbation on the arrhythmogenic potential.

## CONCLUSION

This paper presents a mesoscale soft robotic interface that enables controlled physical perturbation of human organoids with simultaneous functional readout. By integrating compliant actuators with embedded sensing elements^28^, this platform provides a means to directly connect mechanical inputs to tissue-level responses within a single experimental system. Studies with this platform indicate that cortical organoids exhibit distinct responses to trauma, including transient signal silencing, recovery, and cumulative alterations under repeated loading conditions. In cardiac ring tissues, sustained stretch alters spontaneous rhythm stability, conduction dynamics, and repolarization behavior. These results reveal the sensitivity of human organoids to physical perturbations through direct observation of dynamic transitions between functional states under well-defined mechanical conditions.

This framework creates systematic experimental access to force-related biological investigations that are difficult or impossible to perform directly with human tissues due to safety and ethical constraints. By allowing the controlled delivery of defined mechanical inputs, the instrumented soft robotic structures permit quantitative investigation of mechanosensitive signaling and tissue-level responses across previously inaccessible conditions. The technology can be extended to a wide range of mechanically active tissue models, including gastrointestinal, muscle, vascular, lymphatic and multi-tissue assembloid systems, where mechanical forces are essential for physiological regulation and disease progression. Additional possibilities include expanded modes of mechanical control, including rapid actuation, access to high force regimes, and diverse deformation modalities such as shear, cyclic loading, and spatially heterogeneous strain, with potential for improved spatial precision and uniformity of force application. Compatibility with multimodal interrogation, including optical imaging and integrated electrical and mechanical stimulation, will allow direct correlation of mechanical inputs with spatiotemporal functional dynamics. This scalable and broadly applicable approach establishes a foundation for interrogating mechanobiology in human organoids, with potential to uncover force-mediated disease mechanisms and to advance predictive, human-relevant in vitro models of human physiology.

## METHODS

### Nonlinear FEA-based deformation analysis

Numerical simulations were performed using FEA with commercial software (ABAQUS, version 2022). The mass centers of the devices were precisely aligned with the center of the coordinate system in the assembly module. An implicit dynamic step with moderate dissipation was applied, followed by a general static step to eliminate transient effects. General contact was defined in the initial step, employing “hard” normal and frictionless tangential behavior. A pressure load with ramp amplitude was applied to the inner surface of the channel, while the bottom surface of the outer ring was constrained with an encastre boundary condition. The structure was discretized using 8-node 3D stress elements (C3D8RH), with a refined mesh feature size smaller than 1/10 of the channel width. The incompressible second order Yeoh hyper-elastic constitutive model ^24^ was adopted to represent the mechanical properties of the silicone elastomer (Dragon Skin 30, Smooth-on), with the elastic modulus *E_elastomer_* =0.66 MPa, fitted from uniaxial tensile test (*C*_10_=0.11MPa and *C*_01_=5kPa, **Fig S3**). Organoid was modeled as an incompressible Neo-Hooke solid^30^ with the elastic modulus *E*_organoid_=200Pa^31^ (*C*_10_=33Pa). The NOA and PI were modeled as linear elastic materials with moduli *E*_NOA_=1.743GPa^32^ and *E*_PI_=2.5GPa^33^, and Poisson’s ratios *ν*_NOA_=0.49 and *ν*_PI_=0.34^33^, respectively.

### Pneumatic soft robot

The fabrication began with the preparation of patterned molds formed with a photo definable epoxy (SU-8 3050, MicroChem) to define the geometries of the soft elastomeric layers with channel structures. Silicon wafers were sequentially cleaned with acetone, isopropyl alcohol (IPA), and deionized (DI) water. The SU-8 was dispensed onto the wafer and spin-cast at 3000 rpm for 60 s, followed by a resting at 65℃ for 20 min to facilitate stress relaxation prior to soft baking on a hot plate at 95℃ for 20 min. The layer was exposed to ultraviolet light (400 mJ•cm^−2^) through a chromium photomask using mask aligner (SUSS MABA6, SUSS Micro Tec), followed by post-exposure baking (PEB) at 95℃ for 4 mins. Development was performed using an automated developer (Osiris UNIXX S 20 D. SUSS Micro Tec.) for 10 min. The patterned mold was baked on a hot plate at 160℃ for 10 min to relieve residual stresses, to eliminate cracks in the patterned features, and to harden the SU-8. The mold prepared in this manner was silanized in a vacuum desiccator using trichloro (1H, 1H, 2H, 2H-perfluorooctyl) silane (97%, Sigma-Aldrich) for 1h to form a self-assembled monolayer (SAM). A commercial formulation of platinum cured silicone elastomer was mixed at a 1:1 ratio. The channel layer resulted from dispensing this mixture onto the mold, spin-casting at 3000 rpm for 30s, and holding at room temperature for 1 h for leveling. For the bottom layer, the mixture was dispensed onto a SAM-treated glass substrate and spin cast at 4000 rpm for 30s. Both layers were cured overnight in an oven at 75℃. The surface of the cured channel layer and a water-soluble tape were exposed to ultraviolet ozone (UVO) for 2 min and then laminated together, to facilitate bonding by baking at 75℃ for 30 min. The replicated channel elastomer was peeled from the SU-8 mold, and a channel inlet was formed using a mechanical punch. The channel layer and the bottom elastomer on glass were surface-treated using a March reactive ion etcher (March IRE, 370 mTorr, 75W, 25 sccm O_2_, 1min), and then immediately laminated to bond them via overnight curing at 75℃. Subsequently, the water-soluble tape was dissolved by rinsing with flowing DI water, and the device was dried at 75℃ for 1 h.

### Thin-film electronics

Glass substrates were cleaned following the same procedure described above. Poly(methyl methacrylate) (PMMA; 950 PMMA A5, Kayaku Advanced Materials) was spin-cast at 3000 rpm for 40 s and baked on a hot plate at 180℃ for 1 h. A polyimide precursor (PI-2545, HD Microsystems) was then spin-cast at 1000 rpm for 1 min, followed by sequential baking at 120℃ for 2 min. and 150℃ for 2 min to remove residual solvent. Imidization was completed in a vacuum oven at 260℃ for 1 h. A negative photoresist (nLOF 2035, MicroChemicals GmbH) was spin-cast at 5000 rpm for 40s, and patterns were defined using a maskless aligner (MLA 150, Heidelberg Instruments). The patterned resist was immersed in developer (AZ 300 MIF, Michrochemicals GmbH) for 35 s, immersed in DI water for 2 min, and dried. Cr and Au layers (10/200nm) were deposited by electron-beam evaporation, followed by lift off with a stripper solution (AZ 400T stripper, MicroChemicals GmbH) for 12 h to define the electrode traces. An encapsulation layer of polyimide was subsequently deposited using the procedures described above. For patterning the encapsulation layer, a 100 nm thick layer of copper deposited by thermal evaporation (Explorer 14, Denton Vacuum) served as a hard mask. Positive photoresist (S1805, Kayaku Advanced Materials) was spin-cast at 4000 rpm for 40 s, followed by exposure, and development (AZ 917 developer, MicroChemicals GmbH) for 30 s. Immersion in a Cu etchant (PC Copper Etchant 100, Transene) diluted with DI water (1:10) for 2 s removed the exposed Cu, in preparation for oxygen plasma dry etching in a March RIE (200W, 200 mTorr, 25 sccm O2) for 90 min. After removing the remaining Cu, rinsing in DI water and drying in air completed the fabrication.

### Assembly platform

The thin, flexible electronics layer was transferred onto the soft robotics layer by attaching thermal tape, immersing it in DI water for 10 min, and releasing it from the glass substrate. Residual PMMA on the backside was removed with acetone. The surfaces of the robotic elastomer and backside of the electronics were treated with ozone plasma for 2 min. A drop of optical adhesive (NOA61, Norland Products) was applied, and then both layers were aligned under an optical microscope using patterned alignment marks. A PET film and glass slide were placed on top, the stack was clamped to maintain conformal pressure. The assembly was then cured in a UV chamber for 1 h. Afterward, the device was placed on hot plate at 120℃ to release the thermal tape, and the device outline was defined using a laser cutter (LPKF ProtoLaser U4, LPKF Laser & Electronics). To form the pneumatic inlet, a 1.5mm diameter hole was punched in a 5mm thick PDMS block. The PDMS block and device surface were cleaned, plasma treated by exposure to an oxygen reactive ion etch (March RIE, 75W, 370mTorr, 25 sccm O_2_, 1 min), and immediately bonded. The assembly was then heated overnight at 75℃ to strengthen the interface. A PDMS well was attached with adhesive (Kwik-Sil, World Precision Instruments), and the electronic pads were connected to a printed circuit board (PCB) via anisotropic conductive film (ACF) cable. Pt black was electroplated onto the electrodes using a potentiostat (Autolab PGSTAT128N, Metrohm AG) in a three-electrode setup. The plating bath contained 3 wt% chloroplatinic acid and 0.1 wt% lead acetate, with a Pt wire as the counter electrode and Ag/AgCl as the reference electrode. Electrodeposition was performed at −0.1 V for 60 s. Devices were sterilized by washing in IPA, exposing to ultraviolet light for 1 h, and rinsing in ethanol shortly before loading the organoid.

### Pneumatic control system

A custom pneumatic actuation system was developed to independently drive four soft robotic arms. The system was powered by a 24-V DC supply distributed through a master on/off switch. The 24-V rail powered both the proportional valve driver boards and DC–DC power converters (HiLetgo, LM2596), which generated regulated 5-V rails for the pressure sensors and control electronics. Compressed air was supplied from the building air line. Each soft robotic arm was controlled by two miniature proportional valves (Kelly Pneumatics Inc., KPI-VP-20-25-13-V), one for inlet flow and one for outlet (exhaust) flow, enabling bidirectional and continuous pressure regulation. The valves had a 0.013-inch orifice, Viton seals, and operated with inlet pressures up to 25 psi. Each valve was driven by a dedicated proportional valve driver board (Kelly Pneumatics Inc., KPI-DRV-1-24-010), which converted a 0–10 V control input into a current-regulated 0–20 V output for stable valve actuation. Inline pressure sensors (Honeywell, ASDXAVX030PGAA5) were installed in each pneumatic channel and powered from the regulated 5-V rail. Their analog outputs were routed to a microcontroller (Arduino Mega 2560 Rev3), which performed real-time pressure monitoring and closed-loop control.

### Closed-loop PID pressure control

Closed-loop regulation of pneumatic pressure was implemented on the Arduino Mega 2560 using a proportional–integral–derivative (PID) controller. Pressure sensor signals were sampled at 1 kHz via a hardware timer interrupt and digitally filtered with a first-order low-pass filter (2-Hz cutoff) to suppress measurement noise while preserving dynamic response. For each pneumatic channel, the filtered pressure signal was compared with a programmable target pressure waveform, and the PID controller computed a control output that modulated the corresponding inlet and outlet proportional valves. The PID gains were set to *K_P_* = 50, *K_I_* = 15, and *K_D_* = 0.01, with output limits constrained to 0–100% of the valve command range and a control update interval of 5 ms. The PID output was converted into an analog control voltage (0–10 V) using high-resolution hardware PWM signals filtered by RC low-pass networks. These voltages were applied to the driver boards (Kelly Pneumatics), which regulated the current delivered to the proportional valves (0–20 V), thereby controlling airflow into and out of each soft robotic actuator.

### Motion tracking of the soft robotic actuators

Actuator deformation and endpoint trajectory were quantified from side-view videos using a custom MATLAB-based analysis pipeline. After contrast enhancement and adaptive threshold-based segmentation, three characteristic points (anchor, midpoint, endpoint) were manually initialized and refined by local feature detection. Trajectories were determined using a Kalman filter based tracker. Spatial calibration was performed using an object of known length. Curvature was computed for each frame from the osculating circle defined by the three tracked points and converted into physical units. Curvature time series were synchronized with video timestamps and used for subsequent analysis.

To describe the deformation of the soft robotic arm, a model was based on a planar, inextensible trajectory curve with a total length *L*. Assuming constant curvature during actuation, *L*, defined as,

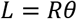

where *R* is the radius of the arc, and *θ* is the bending angle. This configuration yielded cartesian coordinates of trajectory as,

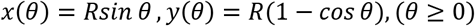

These coordinates were translated into

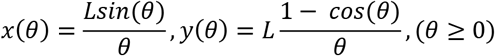

This geometric model was used to predict arm tip trajectories, which were compared with FEA simulations and experimentally tracked motion. The curvature of robotic arms (*κ*) was calculated as the reciprocal of *R*.

### Contact sensor circuits

Contact between the soft robotic arms and the organoid was established by monitoring the impedance between a pair of Au sensing electrodes (each sized at 100 µm × 100 µm and separated by 200 µm) integrated onto each arm. The impedance was quantified by applying an AC excitation voltage to the electrode pair and measuring the resulting AC current. The excitation sinusoidal voltage (100 kHz, 0.4 to 2 Vpp) was produced by a function generator (Rigol DG1032Z) and applied to the electrode pair. The signal was distributed in parallel to the robotic arms to excite all sensing electrodes concurrently using a single function generator output. The AC current between each pair of electrodes was amplified and converted into the voltage domain by a transimpedance amplifier (TIA). Each TIA was implemented using one channel of a quad operational amplifier (TLV3544) and a 100 kΩ feedback resistor. The input of each TIA was virtually grounded to provide a reference for the excitation voltage. The single-supply nature of the operational amplifier was leveraged to rectify the AC current at the TIA input. The output signal of each TIA was averaged using a first-order RC lowpass filter (3.4 Hz cutoff frequency) and routed into the analog input pins of the Arduino Mega 2560. The voltage signal from each robotic arm was sampled at 50 Hz and averaged over 10 samples to reduce the effect of measurement noise. A normalized impedance error signal was generated by comparing the measured voltage to the expected values for the contacted and non-contacted states, obtained through calibration measurements. As the robotic arms were inflated into contact with the organoid, the impedance error signal was used to decrease the rate of change in the pressure setpoint until contact was established.

### Modeling of the contact sensor and impedance-based analysis of the sensitivity

The sensitivity of the contact sensor *S* was defined as

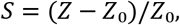

where *Z* and *Z*₀ denote the impedance with and without contact, respectively. Given the significant size mismatch between the electrode (∼140μm) and the organoid (∼1.5mm, not including the surrounding electrolyte), the electrolyte/organoid domain was modeled as a semi-infinite medium. At low frequencies (≤100 kHz), the electrolyte/organoid was approximated as a pure resistance *R_s_*, calculated from finite-element analysis (FEA). The electrode–electrolyte interface was modeled as a constant phase element (CPE) in parallel with a resistance *R_p_*=*kA*^−1^, where *A* is the electrode area and *k* is an interface-dependent constant determined by fitting the experimental data. The total impedance was given by _34_

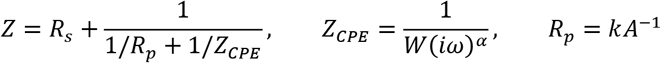

where, *ω* is the angular frequency, and *i* the imaginary unit. The CPE coefficient *W* and exponent *α* were positive constants (smooth, clean surfaces have values of *α* that approach unity, i.e. approaching ideal capacitive behavior _34_) obtained through fitting the experimental data.

When there was no contact, the conductivity of the electrolyte (PBS; *σ*_0_ =1.26S·m⁻¹, range 1.255-1.375S·m⁻¹ _35_) determines the electrolyte resistance *R_s_*_0_ based on FEA. Experimental results for both the frequency and distance *d* were used to fit the parameters as *α*=0.37, *W*=1.7E-7 S·s_α_, and *k*_0_=9.3E-3 Ω·m_2_, as shown in **Fig. S6**. When there was contact, the conductivity of the organoid was modeled as *_36_* 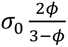, where *φ* is the porosity (range 0.2-0.33 *_37,38_*), *σ*_0_ is the conductivity of electrolyte (PBS, 1.26Sm⁻¹ _35_). A typical value of *φ*=0.27 (in range 0.2-0.33 *37,38*) was used to determine resistance of *Rs* based on FEA.

With the same value of the CPE coefficient *W* and the CPE exponent *α* from the *Z*₀ fitting, the only remaining parameter *k* was determined to be 7.4E-3 Ω·m_2_, as shown in **Fig. S6 G**.

### FEA simulations for the electrolyte/organoid resistance

Numerical simulations were conducted using a commercial software package (COMSOL Multiphysics™). The domain of the electrolyte/organoid was represented as a cubic block with dimensions equal to 45 times the electrode width to approximate a semi-infinite medium. The Electric Current interface within the AC/DC module was employed. A potential of 1 V was applied to the anode, while the cathode was held at zero potential. All remaining boundaries were assigned as electrically insulating. A refined mesh employing tetrahedral elements with feature sizes smaller than 1/5 of the electrode width was adopted to ensure accuracy. A frequency-domain study was conducted to characterize the frequency response, and the resistance was obtained from the reciprocal of the terminal admittance.

### Stem cell cultures and differentiation for cortical organoids

A human induced pluripotent stem cell (hiPSC) line, WTC-11 (Coriell Institute, male; age 30 years at sampling _39_), was used in this study. Cells were maintained in tissue culture-treated 6-well plates (Fisher Scientific, 087721B) coated with Matrigel (Sigma-Aldrich, CLS354277) and maintained in mTeSR Plus medium (StemCell Technologies, 100-0276) supplemented with 1% penicillin-streptomycin (Thermo Fisher Scientific, 15140148). Cultures were incubated at 37 °C with 5% CO₂ and expanded until reaching approximately 70-80% confluency, typically within 4-5 days. For initial cell seeding, 10 µM ROCK inhibitor Y-27632 (Tocris, 1254) was included for the first 20 h to enhance cell survival. Culture medium was refreshed daily. Prior to organoid generation, cells were passaged once as small clumps using Versene solution (Thermo Fisher Scientific, 15040066) upon reaching confluency.

### Generation and preparation of cortical organoids

Human cortical organoids were generated using a dorsal forebrain organoid differentiation kit (StemCell Technologies, 08620) following established protocols adapted from Pasca et al. _40_. WTC-11 cells were dissociated into single-cell suspensions using Gentle Cell Dissociation Reagent (StemCell Technologies, 100-0485). Viable cells were counted using trypan blue (StemCell Technologies, 07050) and a hemocytometer (Fisher Scientific, 0267151B). Approximately 3 × 10⁶ cells were seeded into a single well of an AggreWell 800 plate (StemCell Technologies, 34815) in medium supplemented with 10 µM ROCK inhibitor Y-27632 for the first day. The plate was centrifuged to promote cell capture within the microwells and uniform aggregate formation. From days 1-5, partial media changes were performed daily using medium supplemented with the SMAD pathway inhibitors dorsomorphin and SB-431542 (10 µM each). On day 6, organoids were released from the AggreWell plate, passed through a 37 µm reversible strainer (StemCell Technologies, 27215), and transferred to ultra-low attachment 6-well plates (StemCell Technologies, 38071) at ∼25-35 organoids per well. From days 6-24, complete medium changes were performed every 2 days using medium supplemented with fibroblast growth factor-2 (FGF2) and epidermal growth factor (EGF) (20 ng/mL each) to support neural progenitor expansion. From days 25-43, mitogens were withdrawn and replaced with brain-derived neurotrophic factor (BDNF) and neurotrophin-3 (NT3) (20 ng/mL each) to promote neuronal maturation. From day 43 onward, organoids were maintained in growth factor-free medium with full media changes every 2-3 days. From day 15 until transition to the imaging medium, organoids were treated with a four-compound small-molecule cocktail consisting of GSK2879552 2HCl (Selleckchem, S7796), EPZ-5676 (Selleckchem, S7062), Bay K 8644 (Selleckchem, S7924), and NMDA (Selleckchem, S7072), each at 1 µM, to enhance neuronal maturation, as previously described _41_.

### Viral transduction and calcium imaging

Prior to functional experiments, cortical organoids were transduced with an adeno-associated virus encoding the genetically encoded calcium indicator GCaMP6f (Addgene, 100837-AAV1). Transduction was performed at a multiplicity of infection (MOI) of 5 × 10⁵, followed by incubation for 72 h. Viral particles were then removed, and organoids were transferred to imaging medium consisting of BrainPhys medium without phenol red (StemCell Technologies, 05791) supplemented with 2% B-27 (Thermo Fisher Scientific, 17504044), 1% N-2 (Thermo Fisher Scientific, 17502048), and 1% penicillin–streptomycin, with additional supplements of 1 mM dibutyryl-cAMP (Fisher Scientific, D422825MG), 200 nM ascorbic acid (Sigma-Aldrich, A4403-100MG), 20 ng/mL BDNF (StemCell Technologies, 78005), 20 ng/mL GDNF (StemCell Technologies, 78058), and 10 mM D-glucose (Thermo Fisher, A2494001) to enhance neuronal activity. Organoids were maintained in this imaging medium for at least two weeks prior to experimentation. On the day of the experiment, the imaging medium was refreshed 2-3 h prior to data acquisition. For device insertion and calcium imaging, organoids were maintained in BrainPhys medium without phenol red (StemCell Technologies, 05791) without additional supplements. Organoids were then carefully inserted into the experimental device and placed in a stage-top incubator maintained at 37 °C and mounted on the confocal microscope. To minimize mechanical manipulation-induced stress, organoids were allowed to rest for 10-20 min prior to imaging.

### Mechanical trauma modeling on cortical organoids

For electrophysiological measurements, the electrodes on the soft robotic platform were connected via cables to a commercial recording system (Intan RHD system, Intan Technologies). Electrophysiological signals were acquired using this acquisition hardware and associated software (Intan RHX) at 16-bit resolution and sampling rate of 20 kHz. Time-lapse calcium imaging was performed using an Olympus FV3000 confocal microscope at an acquisition rate of approximately 1 Hz. Electrophysiological recording and calcium imaging were initiated and terminated simultaneously to ensure precise temporal synchronization. During baseline measurements, each organoid was gently gripped using a custom pneumatic valve system, with the applied pressure adjusted according to organoid gripping conditions. Mechanical trauma was delivered as a 1s transient pressure stimulus consisting of a 500 ms pressurization phase followed by a 500 ms release phase. Each experiment included a 3-min baseline recording period prior to mechanical trauma and a 3-min post-trauma period. Control organoids were subjected to identical handling and imaging conditions but did not receive mechanical trauma. For the experiment with three control and three trauma organoids, the organoids were 253 days old at the time of the experiment.

### Immunofluorescence and imaging

Following experiments, organoids were fixed overnight at 4 °C in 4% paraformaldehyde (PFA; Thermo Fisher Scientific, J61899.AK). Fixed organoids were rinsed three times with DPBS (Thermo Fisher Scientific, 14190136) and stored in PBS (Fisher Scientific, 10010031) at 4 °C. For cryoprotection, organoids were incubated in 30% sucrose (Sigma-Aldrich, S0389-500G) diluted in DPBS at 4 °C for 24 h. Organoids were then embedded in OCT compound (Leica Systems, 3801480), frozen, and stored at −80 °C until sectioning. Organoids were cryo-sectioned at a thickness of 10 µm using a cryostat (Microm HM525) and mounted onto Superfrost Plus glass slides (Fisher Scientific, 22-037-246). Slides were stored at −80 °C until immunostaining. Prior to staining, slides were thawed at room temperature and washed with DPBS to remove residual OCT. Sections were blocked for 1 h at room temperature in PBS containing 5% normal donkey serum (Jackson Laboratories, 017-000-121), 3% bovine serum albumin (Thermo Fisher Scientific, 15260037), and 0.3% Triton X-100 (Sigma-Aldrich, X100-5ML). Primary antibodies diluted in blocking solution were applied to sections and incubated overnight at 4 °C. Sections were then washed three times with DPBS and incubated for 1 h at room temperature in the dark with Alexa Fluor–conjugated secondary antibodies diluted in blocking solution. After washing, sections were counterstained with DAPI (1 µg/mL) for 10 min at room temperature, washed again, and air-dried. Slides were cover-slipped using antifade mounting medium (Vector Laboratories, H-1700), protected from light, and stored at 4 °C until imaging. MAP2 and S100B immunostaining was performed to qualitatively confirm neuronal and astrocytic identity. APP and GFAP immunostaining were performed as complementary trauma-associated biomarkers to assess axonal transport disruption and astrocytic reactivity, respectively. Primary antibodies included MAP2 (chicken, 1:5000; Thermo Fisher Scientific, PA1-10005), S100B (mouse, 1:100; Sigma-Aldrich, S2532-100UL), APP (mouse, 1:100; Thermo Fisher Scientific, 13-0200), and GFAP (chicken, 1:500; Aves Labs). Secondary antibodies included Alexa Fluor 488 donkey anti-chicken and Alexa Fluor 647 donkey anti-mouse (1:400; Jackson Laboratories). Sections stained for DAPI, MAP2, and S100B were imaged using a confocal microscope with a 20x objective for qualitative assessment. Sections stained for DAPI, APP, and GFAP were imaged using a 4x objective to obtain whole-section images for subsequent quantitative analysis. Imaging parameters were kept identical across samples stained with the same primary antibodies. Primary-antibody-free control sections were included to confirm signal specificity and ensure appropriate laser power settings.

### Histological image analysis

All image analysis was performed using ImageJ/Fiji. Three-channel fluorescence images were imported using the Bio-Formats importer and processed in batch using a custom ImageJ macro to ensure identical processing across all samples. Uneven background illumination was corrected using rolling-ball background subtraction, with the radius set to a constant value for all images. Sections immunostained for MAP2 and S100B were analyzed qualitatively to confirm neuronal and astrocytic identity and overall tissue organization and were not included in quantitative analysis. Quantitative image analysis was performed exclusively on sections immunostained for DAPI, APP, and GFAP. For quantitative measurements, the DAPI channel was used to define the tissue region of interest (ROI). The DAPI signal was automatically thresholded using Otsu’s method, converted to a binary mask, and used to generate a selection corresponding to the nuclear-defined tissue area. APP and GFAP signals were quantified within the DAPI-defined ROI. For each marker, the corresponding fluorescence channel was independently thresholded using Otsu’s method and converted to a binary mask. Marker-positive area was calculated as the percentage of pixels above threshold within the tissue ROI, providing a normalized measure that accounts for variations in section area. All values were computed directly from raw pixel statistics to avoid dependence on intensity-based measurements.

### Cardiomyocyte differentiation

Human induced pluripotent stem cells (hiPSCs) lines, including patient-derived hiPSC *LMNA* line (gift from Eric Olson, UTSW _29_), were used in this study. Cells were maintained in mTeSR Plus medium (StemCell Technologies, 100-0276) on Matrigel-coated plates (Corning, 354230) and passaged in the presence of 2 µM thiazovivin (TZV; StemCell Technologies, 72254). Cardiomyocyte differentiation was performed using a chemically defined small-molecule protocol involving temporal modulation of Wnt signaling with CHIR99021 and Wnt-C59 _42_. Cardiomyocytes were enriched at approximately day 10 (days 8–12) of differentiation using magnetic-activated cell sorting (MACS)_43_. Following purification, cells were resuspended in RPMI 1640 supplemented with B-27 supplement (Thermo Fisher Scientific, 17504-044), 10% KnockOut Serum Replacement (KOSR; Thermo Fisher Scientific), and 2 µM TZV prior to counting. Cardiomyocytes were subsequently expanded in T75 flasks in the presence of 2 µM CHIR99021 for 11 days _44_. Human cardiac fibroblasts (HCFs; PromoCell, C-12375) were thawed from cryopreserved stocks and recovered in the same medium prior to use.

### Generation of fibrin-based engineered heart tissues

Engineered heart tissues (EHTs) were generated using a fibrin-based hydrogel protocol adapted from prior publications_27_. Cardiomyocytes and HCFs were combined at a 90:10 ratio and pelleted by centrifugation (200 × g, 5 min, room temperature). Fibrinogen from bovine plasma (Sigma-Aldrich, F8630) was dissolved at 10 mg/mL in 10% horse serum medium and sterile-filtered through a 0.2 µm syringe filter. An extracellular matrix mixture consisting of 2× RPMI 1640 (Thermo Fisher Scientific, 31800022), fibrinogen solution, and Matrigel was prepared at a 40:40:20 volume ratio on ice. The cell pellet was resuspended in the matrix mixture prior to seeding. PDMS molds were coated overnight at 4 °C with 5% Pluronic F-127 (Sigma-Aldrich, P2443-250G) in PBS. Thrombin from human plasma (Sigma-Aldrich, T6884) was added at 0.2 U per mg fibrinogen, and the mixture was seeded into the molds (75 µL per well for 6-mm molds; 62.5 µL per well for 5-mm molds). Constructs were incubated at 37 °C for 5 min. Culture medium consisting of low-glucose Dulbecco’s Modified Eagle Medium with HEPES (Thermo, 12320032) supplemented with 10% horse serum (New Zealand origin; (Thermo Fisher Scientific, 26050088), 1% penicillin-streptomycin (Thermo Fisher Scientific, 15070-063), and 0.1% human insulin solution (Sigma Aldrich, I9278-5ML) was added following gelation. For the first 3 days, medium was supplemented with transforming growth factor-β1 (5 ng/mL; PeproTech, 100-21C). Thereafter, tissues were maintained in the same medium without TGF-β1, with medium exchanges performed three times per week.

### Stretching perturbation on EHTs

The soft robotic platform for EHTs was connected to a commercial recording system (Intan RHS system, Intan Technologies). Field potential signals were acquired as described above. Calcium and voltage imaging were performed using a dual optical imaging system using CMOS cameras (MiCAM, SciMedia), described in our previous paper _45_. The platform was maintained on a temperature-controlled heating pad at 37 °C for the duration of the experiment. EHTs were incubated in Tyrode’s solution containing 15µM blebbistatin (Cayman Chemicals, 13186) for 10 minutes. The tissues were then stained with a calcium indicator dye (Rhod-2 AM; 30 µL of 1 mg/ml dye stock solution, <u>Thermo</u> Fisher Scientific R1244 + 30 μl Pluronic F-127 Biotium, 59005), voltage indicator dye (RH237; 30 μl of 1.25 mg/ml dye stock solution, Biotium 61018) together in Tyrode’s solution (dyes and Pluronic combined with 940µL Tyrode’s solution) for 1 min prior to mounting onto the device. Ring-shaped tissues were positioned on the platform and actuated using a PID-controlled pneumatic valve system to establish stable contact with the integrated electrodes. Baseline calcium imaging and electrophysiological recordings were acquired during biphasic electrical pacing (400 µs pulse duration, 1 mA amplitude, 1–1.75 Hz frequency). Mechanical stretching perturbation was induced by gradually increasing pneumatic pressure over approximately 10 s. Following stretching, voltage and calcium imaging and electrophysiological recordings were repeated under the same pacing conditions.

### Electrophysiology signal processing

Electrophysiological dataset used in this work was acquired through RHS head stage and controller (INTAN technology, USA). A customized MATLAB code was used in post-processing this dataset, including zero-phase notch, bandpass filters and discrete wavelet transform for denoising. For WTC-11, ECG-like LFP was obtained by bandpass filter which has a window from 0.1 Hz to 250 Hz, and another window, from 300 Hz to 3000 Hz yielded high frequency domain signals. Also, another code served as a pipeline producing spike sort and raster plot results. For hiPSC-LMNA, high-pass filter (0.1 Hz) was applied to remove baseline wander.

### Analysis of fluorescence dynamics

Fluorescence videos (512 × 512 pixels, 7 fps) were analyzed using custom MATLAB scripts. The green channel of the first frame was extracted and smoothed using a Gaussian filter (σ = 2). Candidate fluorescent spots were identified by thresholding followed by connected-component analysis, and the brightest regions were selected as regions of interest (ROIs) based on their mean intensities. Within each ROI, spot positions were estimated using threshold-based weighted centroid analysis, in which pixels exceeding a fraction (k) of the 99.5th percentile intensity were selected to compute the intensity-weighted centroid. Fluorescence intensity was quantified using a local signal–background subtraction method. For each tracked position, a circular signal region (*R*=15 pixels) and an annular background region (*R_in_*=18 pixels, *R_out_*=28 pixels) were defined. The mean background intensity was subtracted from the mean signal intensity to obtain the net fluorescence intensity,

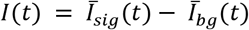

where *I̅_sig_*(*t*) and *I̅_bg_*(*t*) denote the mean pixel intensities within the signal and background regions, respectively.

In addition to intensity-based measurements, area-based fluorescence metrics were computed within the tracking ROI. The area metric *A*(*t*), was defined as the number of pixels exceeding the adaptive threshold. The integrated fluorescence signal, *S*(*t*), was calculated as the sum of pixel intensities above the threshold,

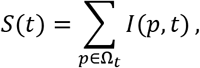

where Ω*_t_* denotes the set of ROI pixels (*p*) whose intensities exceed the adaptive threshold at time *t*. These metrics provide complementary information on the spatial extent and integrated brightness of fluorescence spots. Adaptive threshold parameters were optimized for each dataset using an automated grid-search procedure. Two thresholding strategies were evaluated: percentile-based thresholding (*T* = *k* ⋅ *P*_99.5_)and background-based thresholding (*T* = *μ_bg_* + *ασ_bg_*). Candidate values of *k*(0.35–0.85) and *α*(1.0–4.5) were tested using the first 250 frames of each video. Each parameter set was evaluated using a consistency score incorporating (i) the correlation between *I*(*t*)and *S*(*t*), (ii) the fraction of frames with valid *A*(*t*)values, and (iii) a penalty for high-frequency fluctuations in *A*(*t*). The parameter combination yielding the highest score was selected for full-length analysis. All data processing and visualization were conducted in MATLAB.

## ACKNOWLEDGMENTS

Research reported in this publication was supported by the National Institute of Neurological Disorders and Stroke (NINDS) of the National Institutes of Health under grant number R01NS113935; the National Heart, Lung, and Blood Institute (NHLBI) under grant numbers R33HL168758 and K08HL163392; and the National Institute of Biomedical Imaging and Bioengineering (NIBIB) under grant number R41EB037484. Fifty percent of the research project was financed with federal funds, and 50% was financed with non-governmental funds. In addition, this work was supported by the Querrey-Simpson Institute for Bioelectronics at Northwestern University and by grants R41EB037484 and R01NS113935 (J.D.F.) and K08HL163392 (D.E.F.) from the National Institutes of Health. We thank Serkan Butun for his assistance with microfabrication. This work made use of the Micro/Nano Fabrication Facility (NUFAB) of Northwestern University’s Atomic and Nanoscale Characterization Experimental (NUANCE) Center.

## AUTHOR CONTRIBUTIONS

Conceptualization, *W.-Y.M and J.A.R*; Methodology of soft robotics, *W-Y.M, Z.L, K.S.K, L.P.P and S.L;* Methodology of control, *K.H.H, V.L, Y.M, S.X;* Methodology of biology, *W.-Y.M, Y.S, I.A.C, B.L,C. W.H,N.L, E.M.M, I.R.E, D.E.F and J.D.F;* Data Analysis, *W.-Y.M, K.S.K;* Investigation, *W-Y.M, Z.L, Y.S, K.H.H, K.S.K, T.W.P, Y.J.J, H.Y.A, S.Y, S.O, J.K, D.E.F;* writing – original draft, *W.-Y.M;* writing – review & editing, *Z.L, Y.S, K.H.H, K.S.K E.M.M, I.R.E, D.E.F, Y.H, J.D.F and J.A.R.;* funding acquisition, *D.E.F, J.D.F and J.A.R;*Resources, *J.A.R;* supervision, *Y.H, J.D.F and J.A.R*.

## RESOURCES TABLE

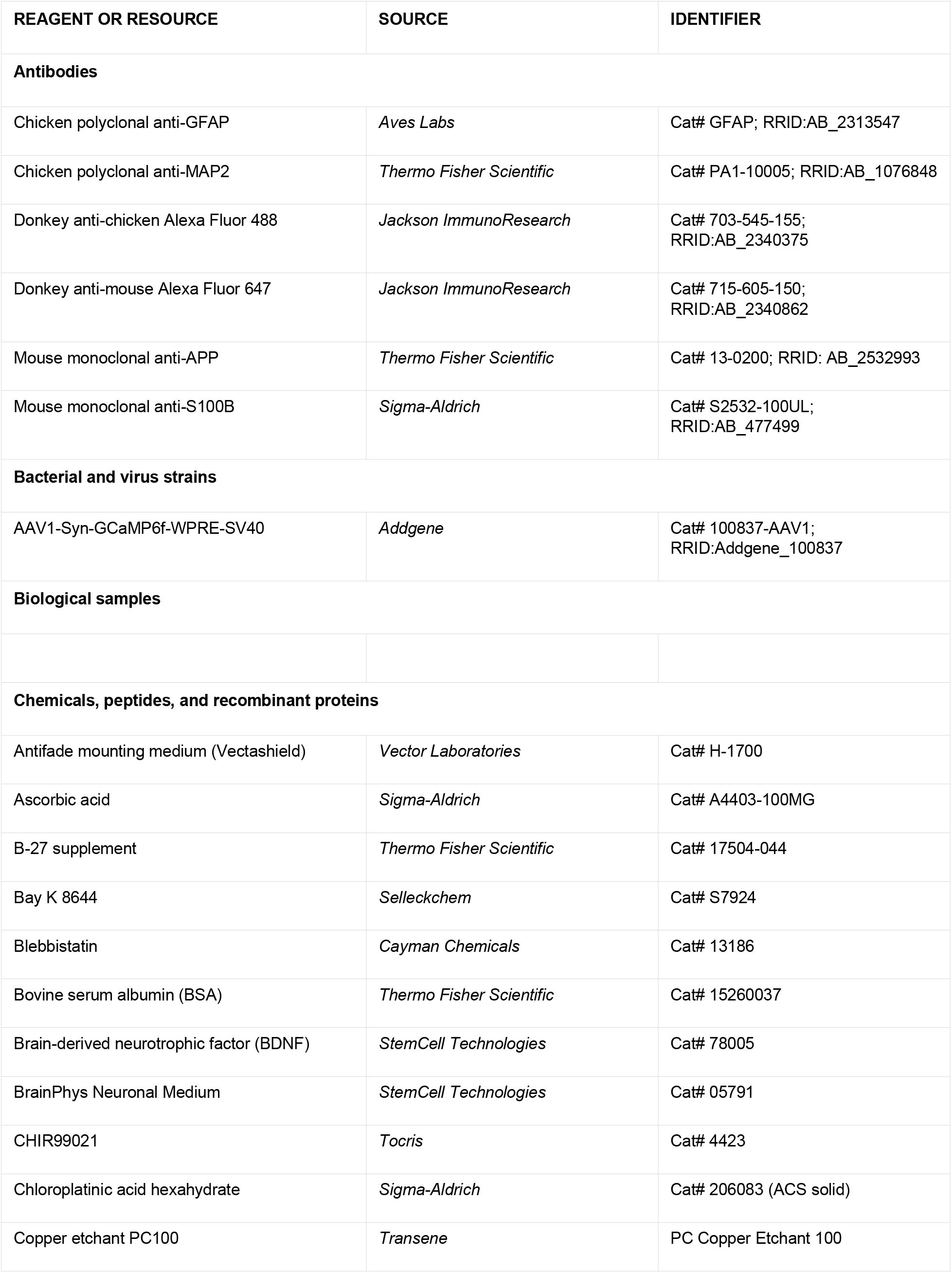
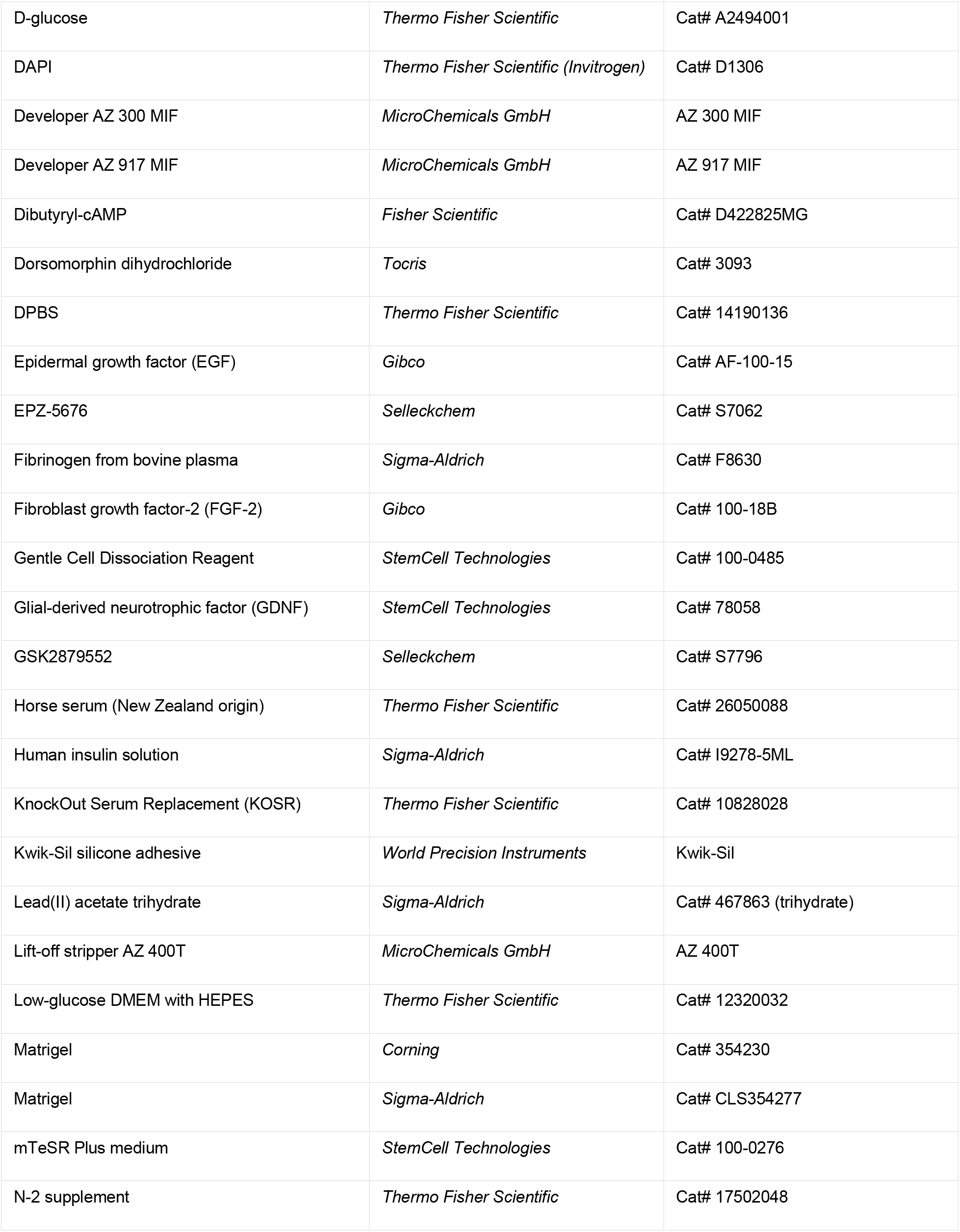
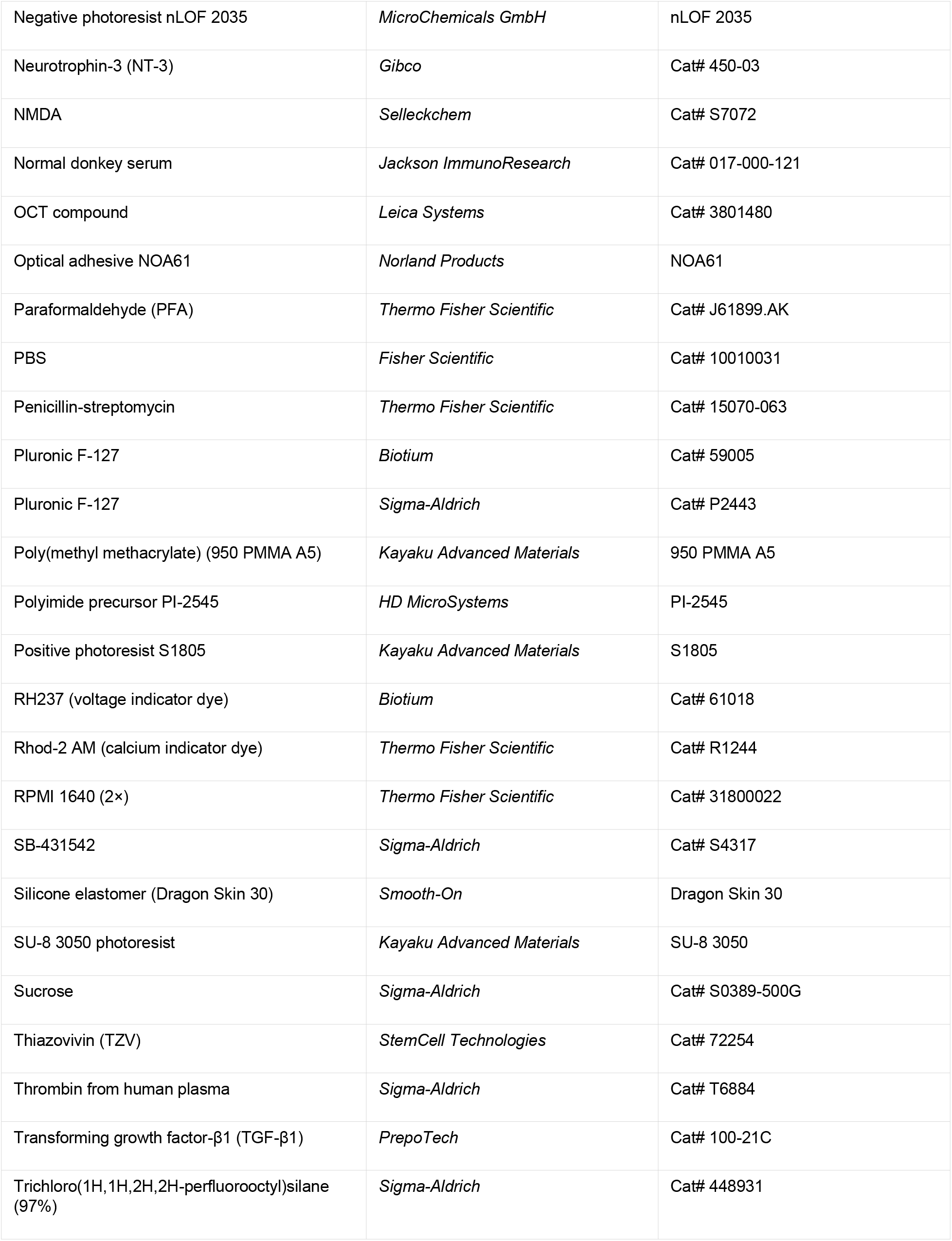
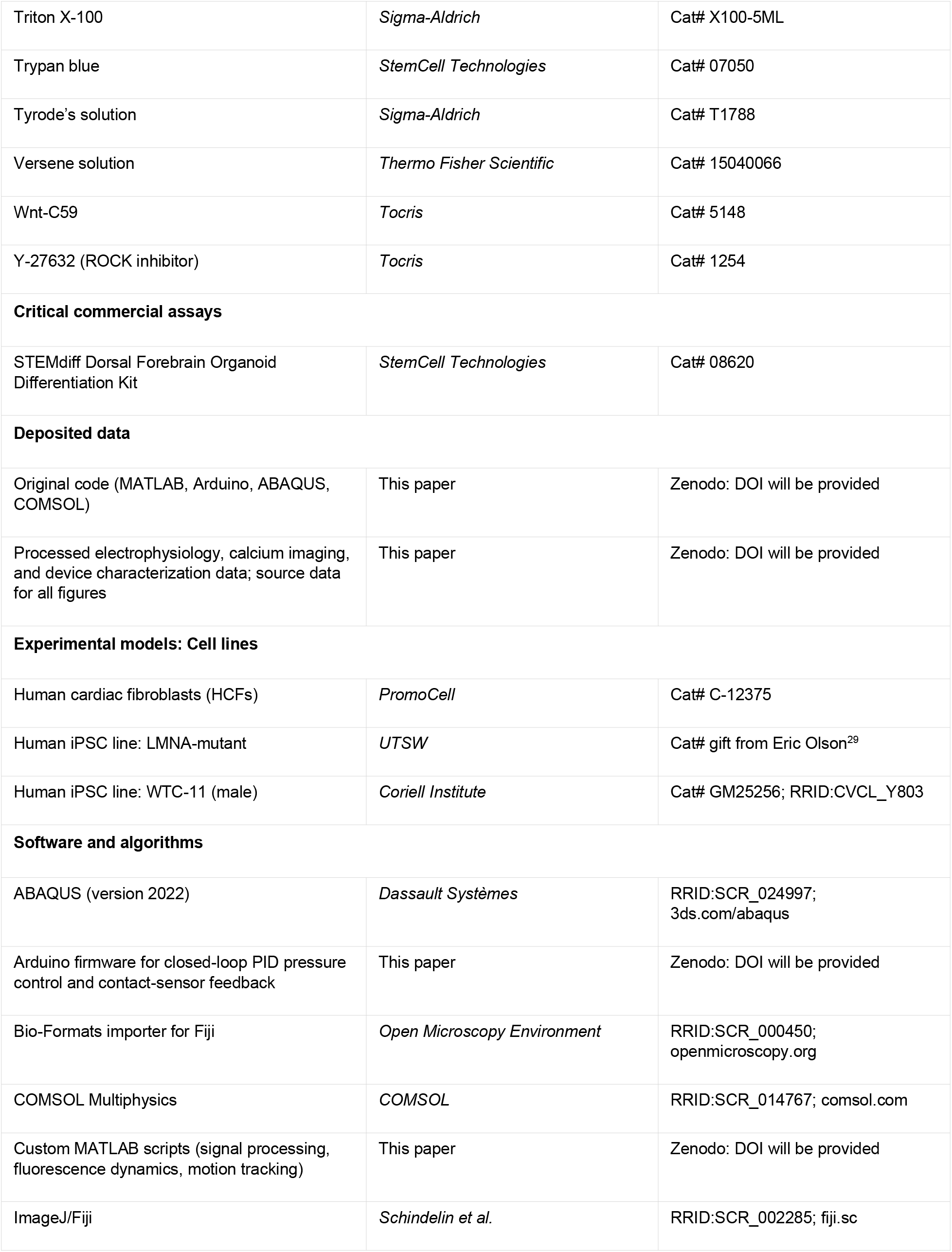
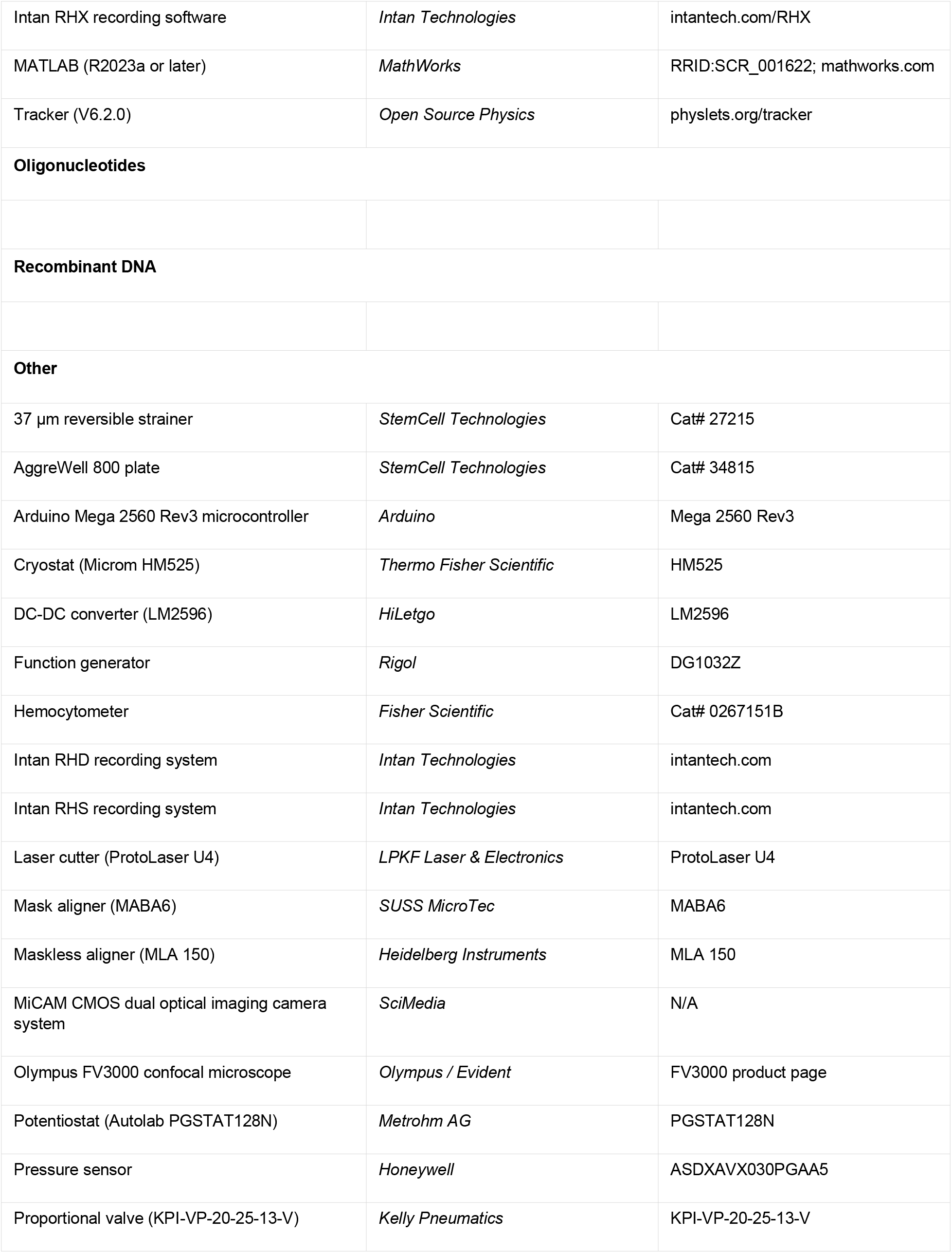
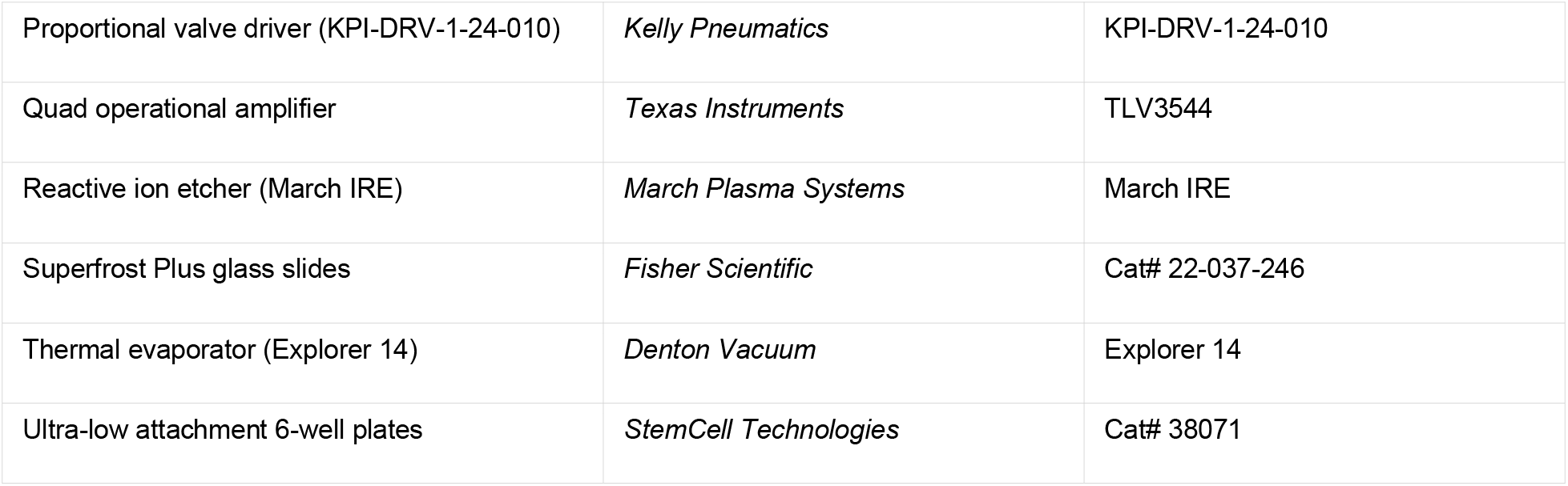

