## Supplemental Information for "Mesoscale Soft Robotics for Mechanically Perturbing and Functionally Monitoring Responses in Human Organoids"

[1]Querrey Simpson Institute for Bioelectronics, Northwestern University, Evanston, IL, 60208, USA.
[2]Center for Bio-Integrated Electronics, Northwestern University, Evanston, IL, 60208, USA.
[3]Department of Civil and Environmental Engineering, Northwestern University, Evanston, IL, USA
[4]Department of Mechanical and Industrial Engineering, University of Illinois Chicago, Chicago, IL, USA
[5]Department of Bio and Brain Engineering, Korea Advanced Institute of Science and Technology, Daejeon, 34141, Republic of Korea
[6]School of Biomedical Engineering, Korea University, Seoul, 02841, Republic of Korea.
[7]Interdisciplinary Program in Precision Public Health, Korea University, Seoul, 02841, Republic of Korea.
[8]Department of Chemistry, Vellore Institute of Technology, Vellore, Tamil Nadu 632014, India
[9]Center for Genetic Medicine, Bluhm Cardiovascular Institute, Northwestern University Feinberg School of Medicine, Chicago, IL, 60611, USA.
[10]Department of Biomedical Engineering, Northwestern University, Evanston, IL, 60208, USA.
[11]Department of Mechanical Science and Engineering, University of Illinois Urbana-Champaign, Urbana, IL, 61801,USA
[12]Department of Mechanical Engineering, Vanderbilt University, Nashville, TN, 37240, USA
[13]School of Glocal Leaders, Glocal Honors College, Inje University, Gimhae 50834, Republic of Korea.
[14]Division of Electrical Engineering, Hanyang University ERICA, Ansan, 15588, Republic of Korea.
[15]Department of Mechanical Engineering, Northwestern University, Evanston, IL, USA
[16]Department of Materials Science and Engineering, Northwestern University, Evanston, IL, 60208, USA.
[17]Department of Neurological Surgery, Northwestern University, Chicago, IL, 60611, USA.
[18]These authors contributed equally
[19]Lead Contact

### SUPPLEMENTAL FIGURES

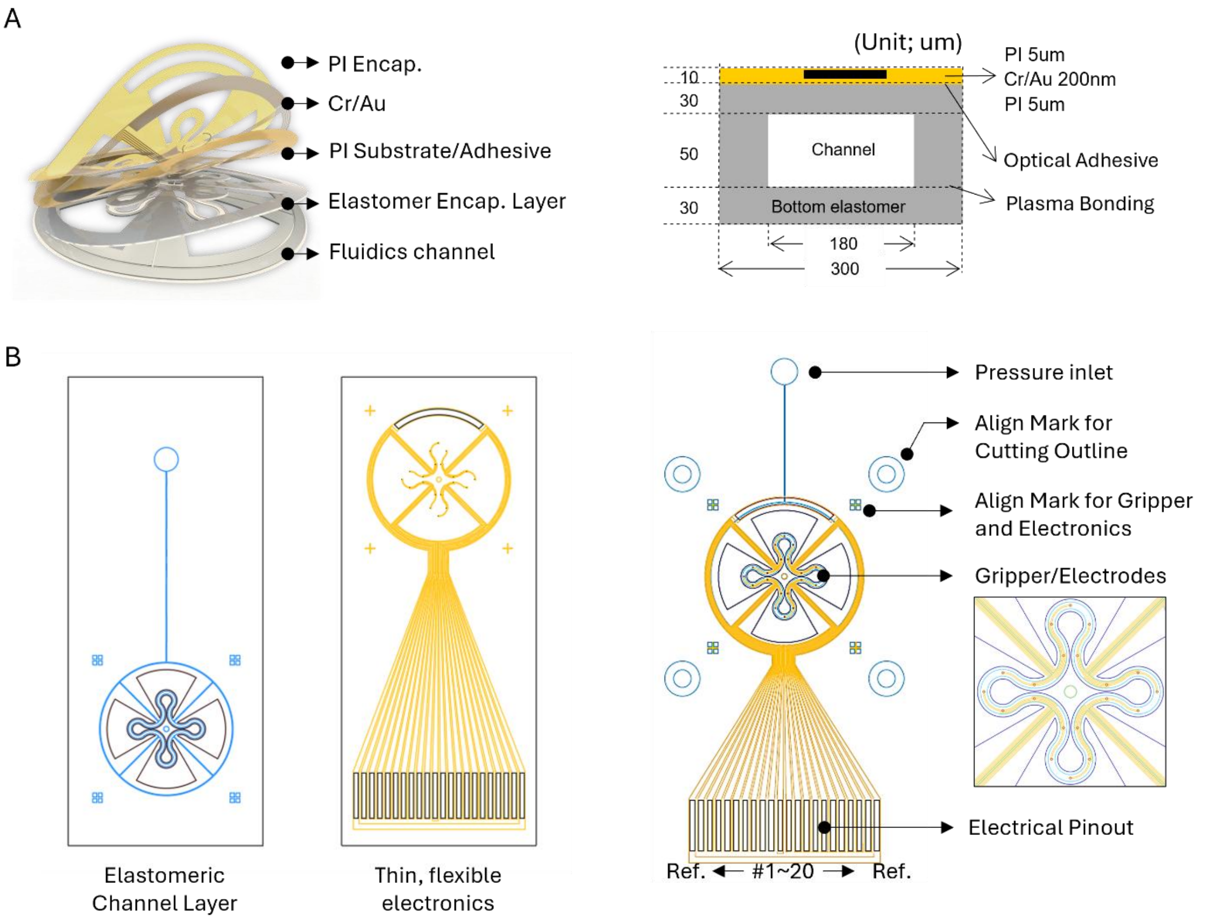


**Figure S1. Layout of the SROI. (A)** Multilayer design of the soft elastomeric channel layer and thin-film electronics, with layer thickness and dimensions. **(B)** Designs of the elastomeric channel layer for soft robotic actuation and the electronic layer for multifunctional integration, shown with an aligned 2D assembled layout.

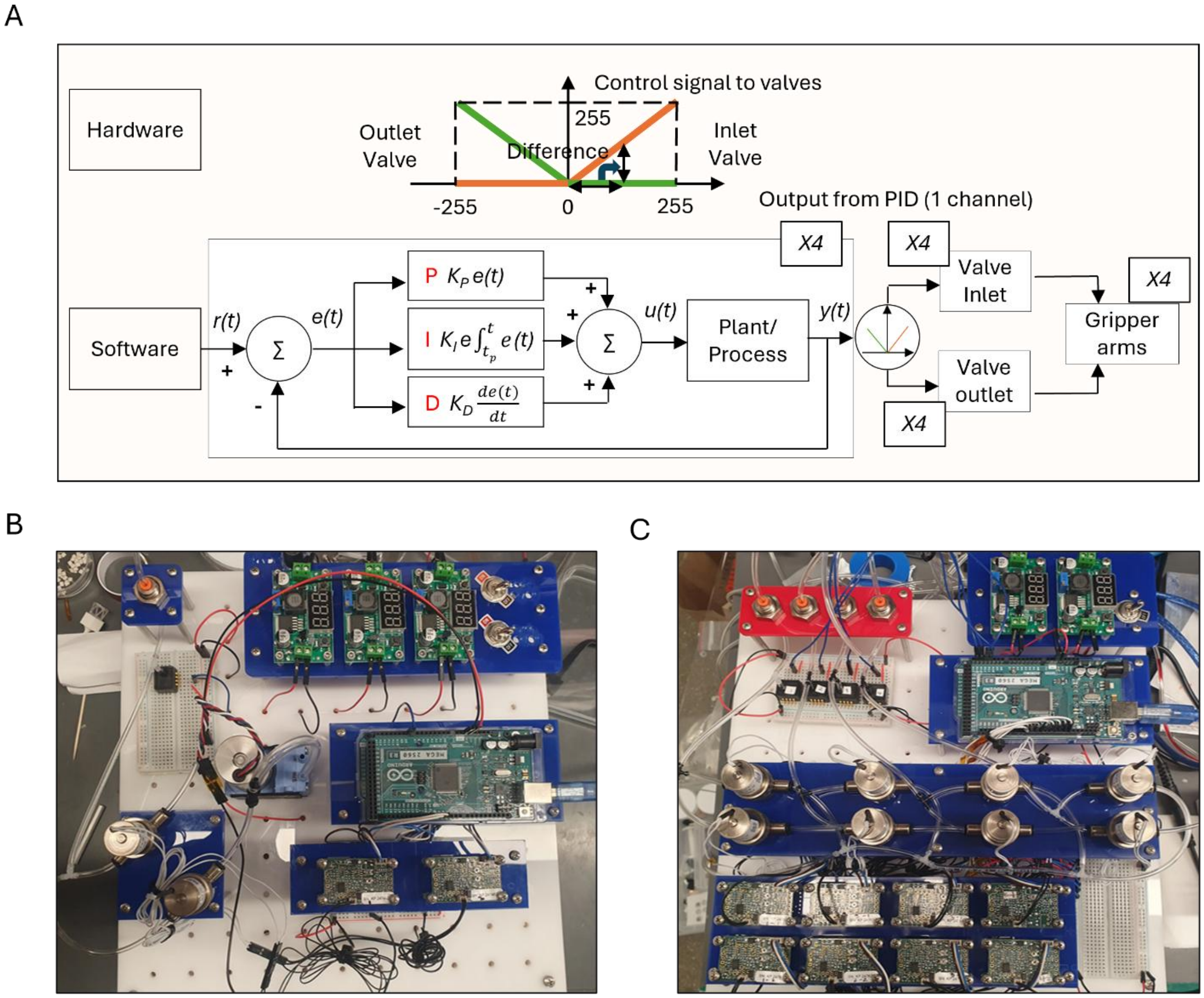


**Figure S2. Custom PID-controlled pneumatic valve system for SROI actuation. (A)** Schematic of the Arduino-based PID control system, illustrating software-hardware integration for differential control of inlet and outlet pneumatic valves. **(B)** Single-channel PID-controlled pneumatic valve hardware. **(C)** Four-channel expanded system for independent control of four gripper arms.

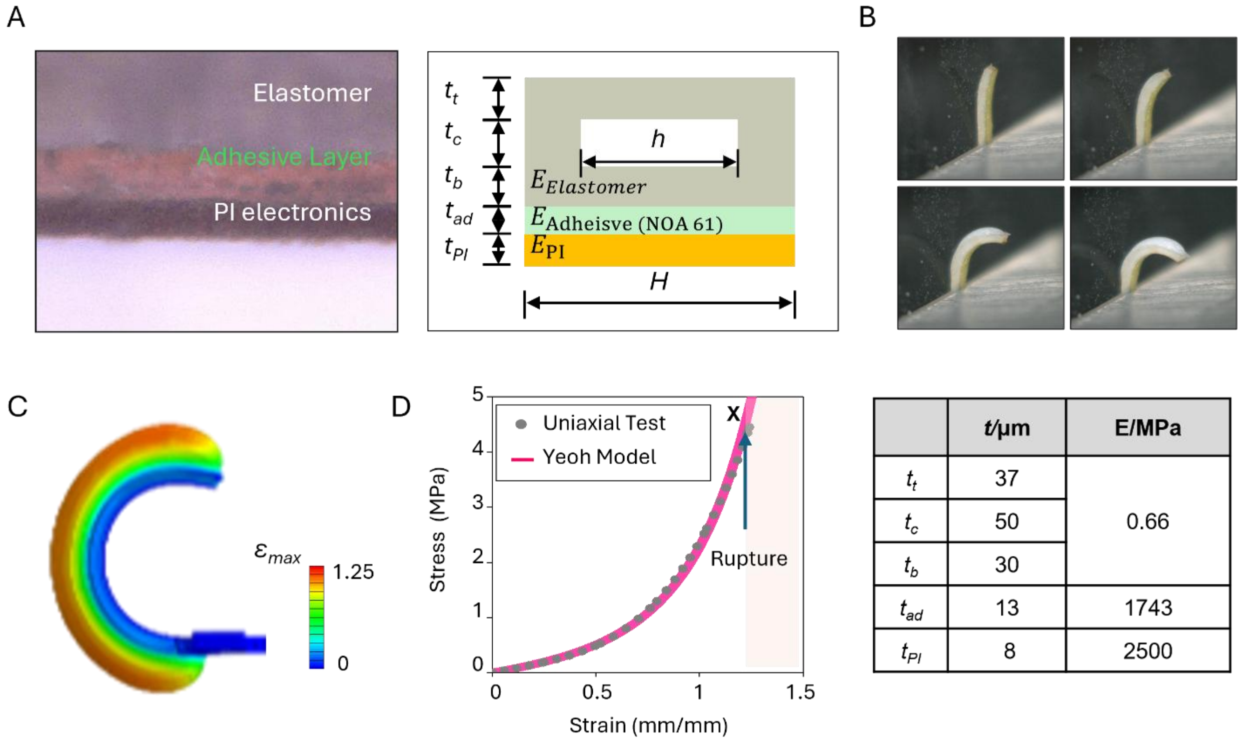


| | $t$/µm | E/MPa |
|---|---|---|
| $t_t$ | 37 | 0.66 |
| $t_c$ | 50 | |
| $t_b$ | 30 | |
| $t_{ad}$ | 13 | 1743 |
| $t_{PI}$ | 8 | 2500 |

**Figure S3. Multilayer interface and bending mechanics of the SROI. (A)** Cross sectional optical micrograph and schematic illustration of a soft robotic grippers and its components. **(B)** Time-lapse bending profiles of a single arm during pneumatic actuation, showing the variation in curvature during actuation. **(C)** FEA of a single arm during pressurization, with the resulting strain distribution. **(D)** Uniaxial tensile testing of the elastomer used in the device (n=10) with a Yeoh hyperelastic fit. The fitted model parameters used for the simulations are reported in the Table.

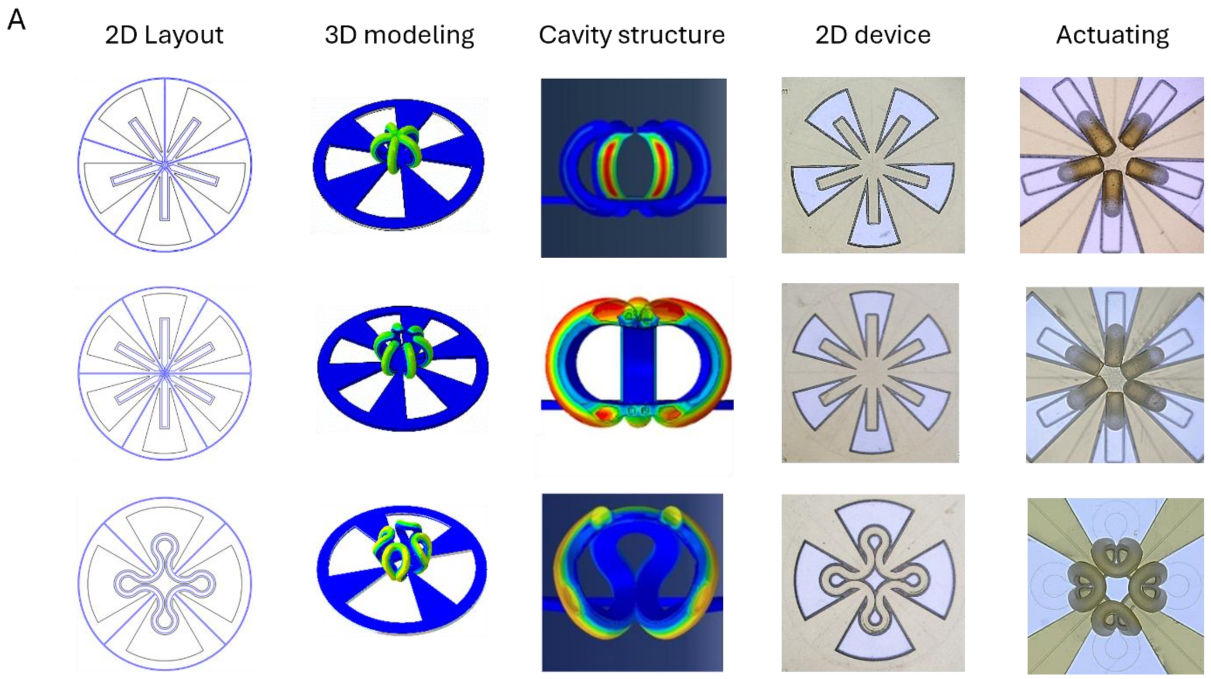


**Figure S4. Additional SROI gripper designs and actuation behavior.** 2D layouts and 3D models of alternative gripper geometries, including 5 and 6 arm designs and 4 arm petal designs. Simulated inner-cavity shapes under pneumatic actuation are shown alongside optical images of the corresponding fabricated devices and their actuated configurations.

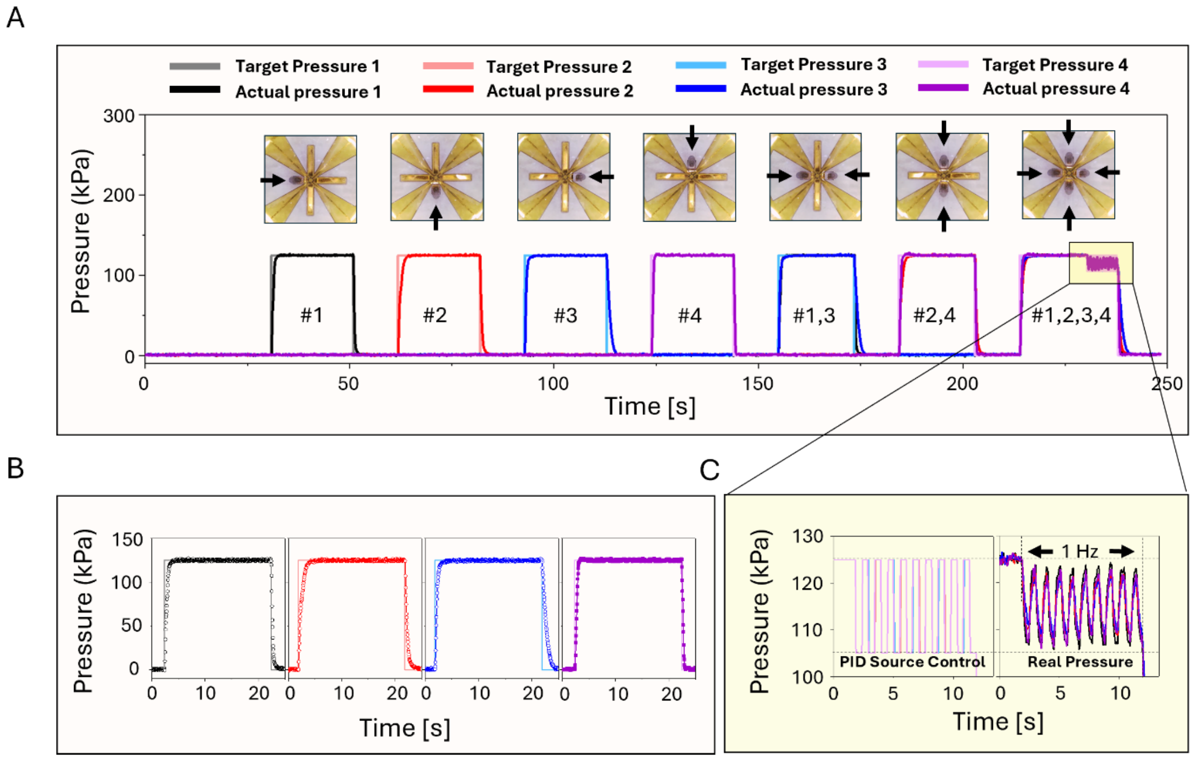


**Figure S5. Independent four-arm actuation with a four-channel pneumatic control system. (A)** Optical snapshots and pressure traces from a four-arm soft robotic gripper driven by a four-channel PID valve system. The sequence includes single-arm actuation (1, #2, #3, #4), paired actuation of opposing arms (#1, 3 and #2, 4), and simultaneous actuation of all four arms (#1-4). Plots show the target and actual channel pressures during the sequence. **(B)** Target versus measured pressures for each channel, demonstrating rapid responses and stable steady-state regulation. **(C)** 1 Hz operation showing PID control of cyclic pressure commands and the corresponding measured pressure response.

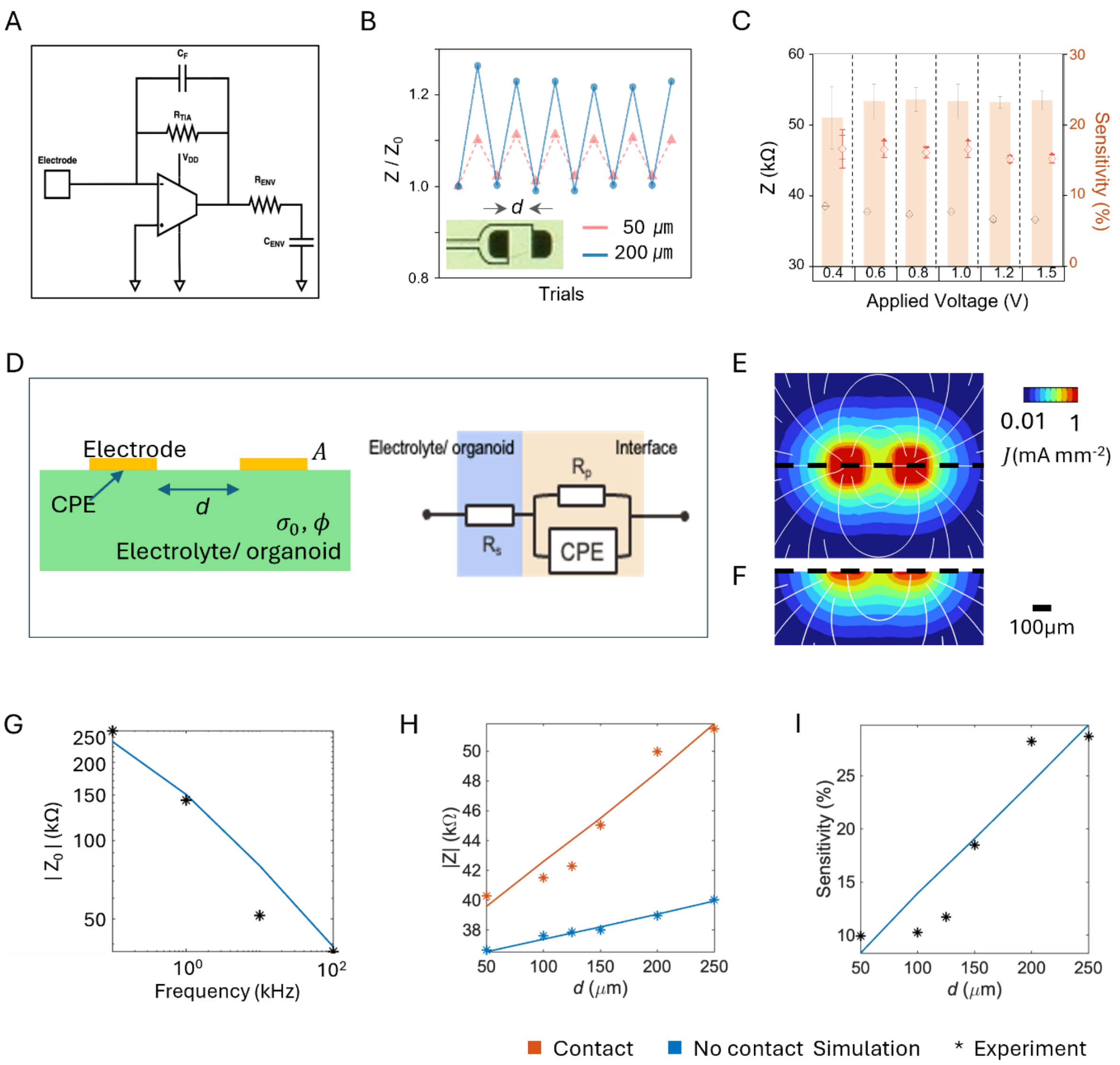


**Figure S6. Impedance-based contact sensing and model validation. (A)** Circuit diagram of the impedance-based contact-sensing readout using a transimpedance amplifier. **(B)** Representative impedance responses for non-contact and contact states measured with two electrode spacing (50 μm and 200 μm). **(C)** Measured impedance changes and contact sensitivity as a function of the applied drive voltage. **(D)** Conceptual schematic illustration of the impedance-based model, the equivalent circuit and the expression for the total inner-electrode impedance. Top **(E)** and side **(F)** views of the internal current density magnitude within the organoid (inter-electrode spacing: 200μm; 2 Vpp, 100 kHz). Solid white lines indicate the direction of the electric field, and black dashed lines denote the cutting plane. The color scale is logarithmic. (G) Frequency responses of total impedance in the non-contact condition. (H) Relation between the total impedance and the electrode distance with and without contact. (I) Relation between sensitivity and the electrode distance.

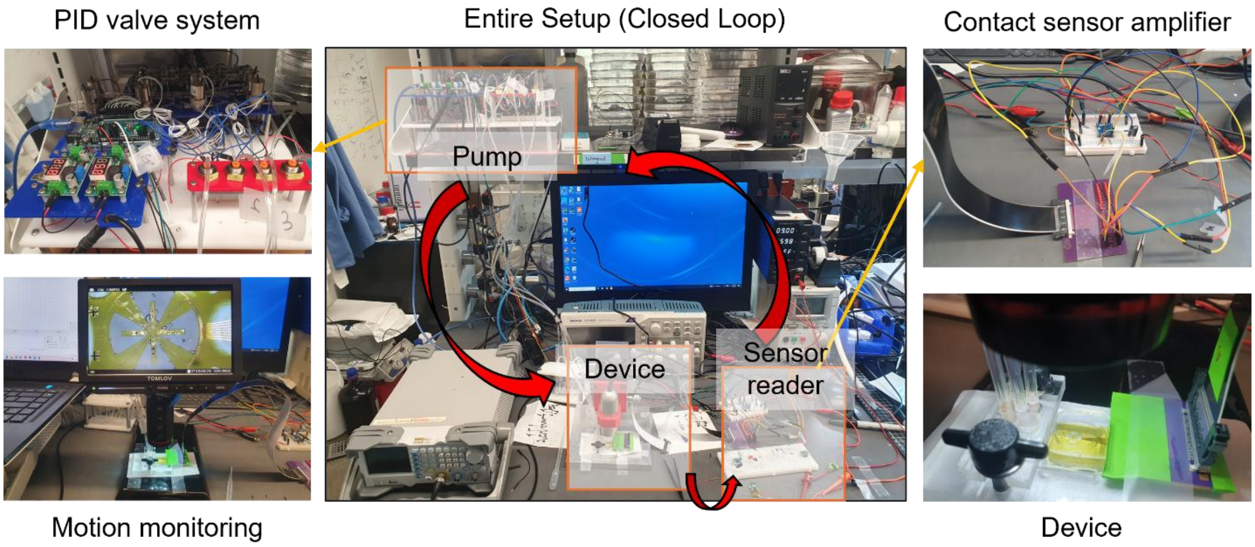


**Figure S7. Closed-loop pressure control system with integrated contact sensing.** Optical images of the full benchtop system integrating (i) an optical microscope for monitoring SROI motion, (ii) a PID-controlled pneumatic valve system connected to the device inlet, and (iii) a contact-sensor readout circuit interfaced with the device electrical connector. The setup enables closed-loop regulation by coupling pneumatic actuation with real-time contact states.

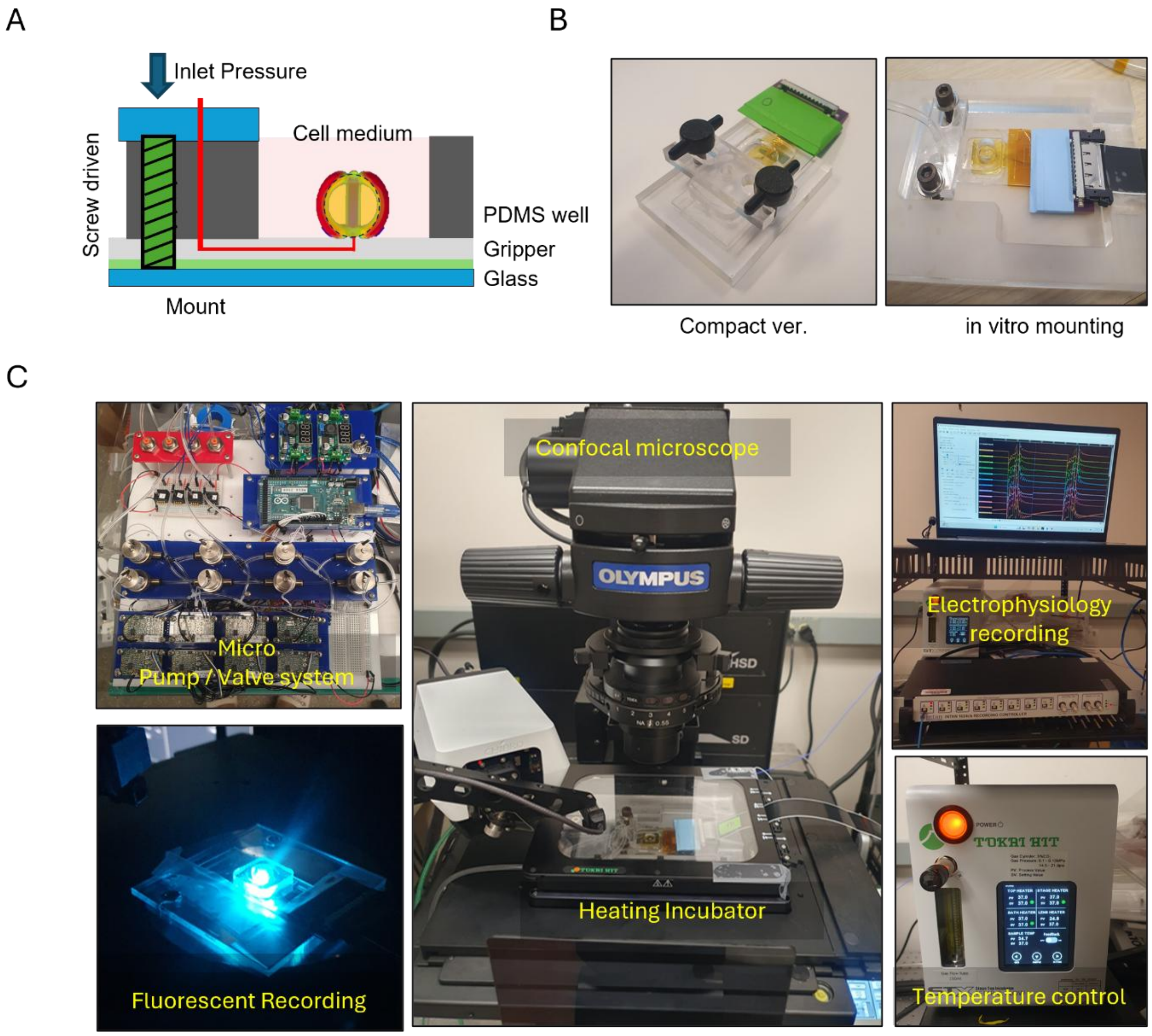


**Figure S8. *In vitro* experimental setup for neural organoid trauma modeling. (A)** Schematic diagram of the assembled device with an integrated reservoir for organoid and culture medium. **(B)** Images of the device mounted on two fixtures; a compact mount (left) and a custom mount designed to integrate with a mini incubator. **(C)** Fully integrated confocal imaging and electrophysiology setup, showing the device positioned in the heated incubator and connected to the pneumatic valve system, MEA recording hardware, and the incubator temperature controller.

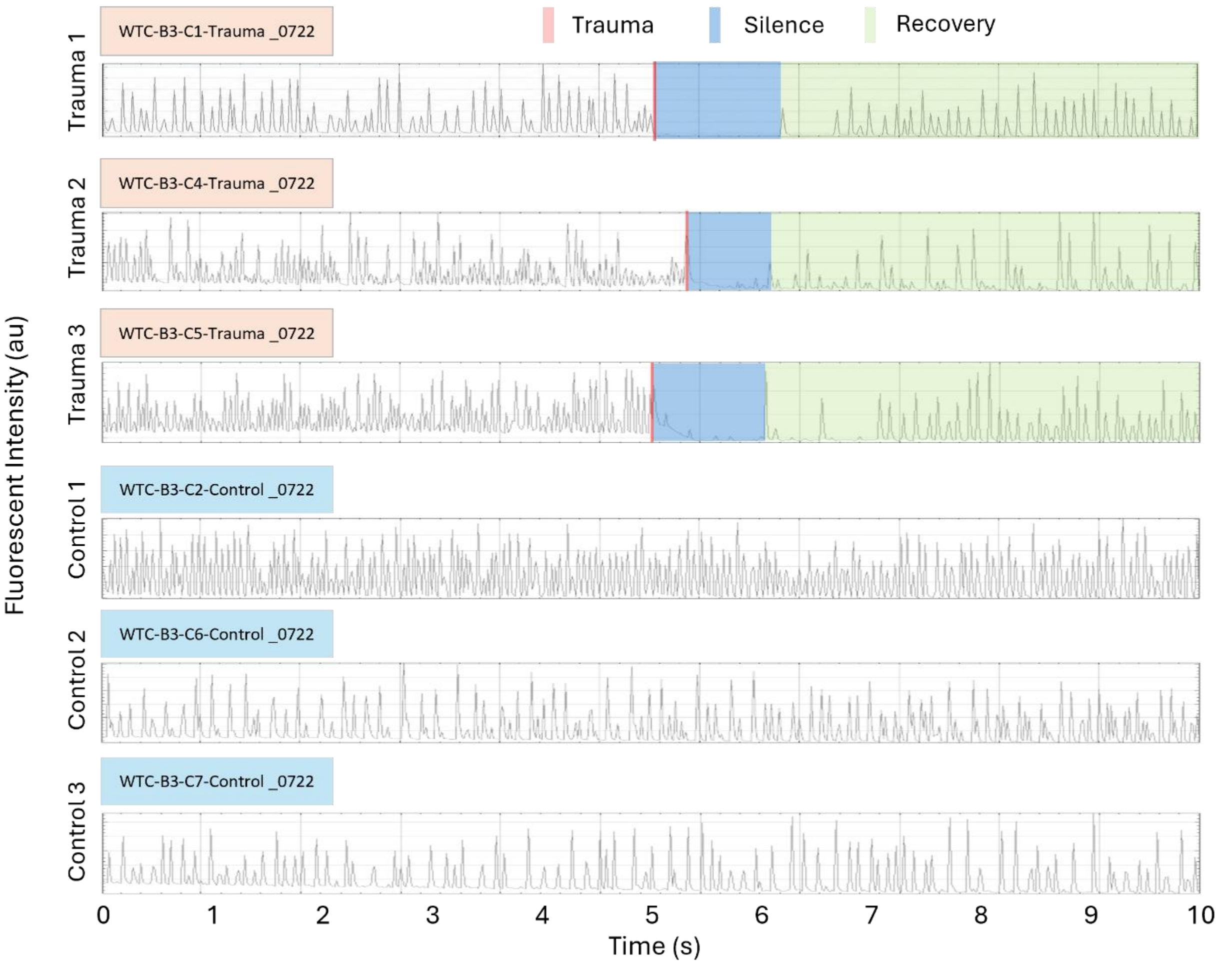


**Figure S9. Reproducibility of trauma-induced Ca2+ responses in neural organoids.** Representative 10-min $Ca^{2+}$ intensity recordings from three independent organoids and three control organoids. In the trauma group, a 1s traumatic loading induces transient silencing of $Ca^{2+}$ activity, followed by a recovery phase. This silencing-recovery pattern is consistently observed across all trauma organoids, whereas control organoids maintain stable $Ca^{2+}$ activity without silencing throughout the recording period.

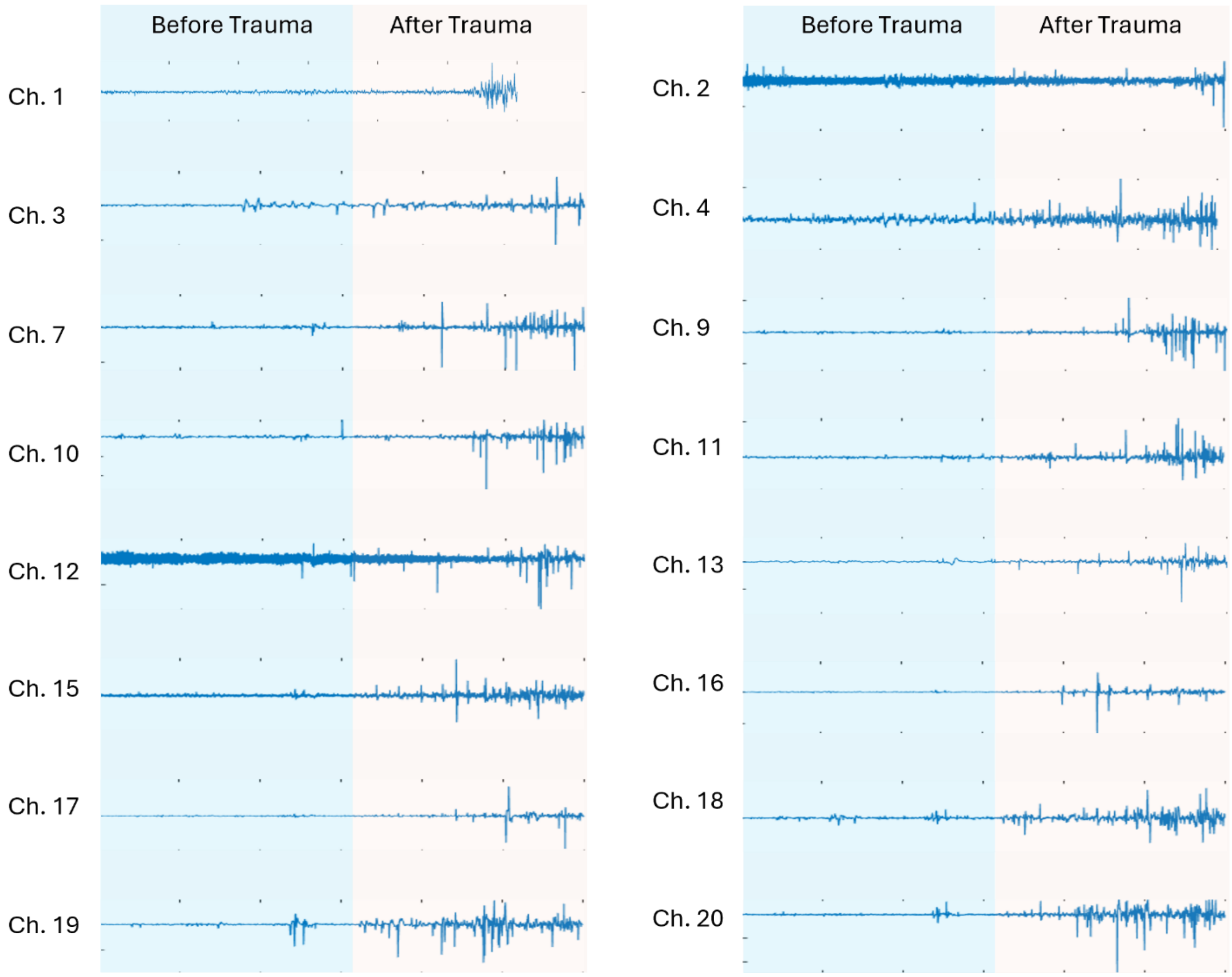


**Figure S10.** In same experiment as Fig. 4D, LFP recordings are relatively quiet before trauma, whereas bursting-like signal appears consistently across 16 electrode channels after trauma.

**SUPPLEMENTAL TEXT**

Each robotic arm incorporates a pair of selectively exposed electrodes (100μm x 100 μm) with an edge-to-edge spacing of 200 μm, as the basis of an impedance-based contact sensor. The sensing scheme employs high-frequency AC excitation (100 kHz, 0.4-2.0 Vpp), in which contact-dependent changes in impedance convert into voltage signals using a transimpedance amplifier (TIA) circuit **(Fig 3D)**. A function generator drives one electrode, and the opposing electrode connects to the TIA input held at virtual ground (0 V). The amplified output voltage differs between non-contact and contact states, yielding a ~20% change upon contact **(Fig 3E)**. Increasing the operating frequency enhances the sensitivity to contact-induced impedance changes **(Fig. 3F)**. Increasing the inter-electrode spacing (50 to 250 μm) increases both the amplitude of the voltage response and the sensitivity, reflecting a larger impedance difference between contact and non-contact states **(Fig. 3G, Fig. S6 D-G)**. The output voltage response exhibits negligible dependence on the applied drive voltage in the range of 0.4 to 2.0 Vpp **(Fig. S6 C)**. A simple equivalent circuit model reproduces these trends. Without contact, the inter-electrode impedance decreases from 240 to 39 kΩ as the frequency increases from 0.1 to 100 kHz, and increases from 37 to 40 kΩ as electrode spacing increases from 50 to 250 μm. By contrast, the contact sensitivity increases from 8 to 30 % over the same spacing range **(Fig. S6 E-G)**. To minimize any possible electrical perturbation to the organoid, operation involves the lowest drive voltage (0.4 Vpp) that preserves a distinguishable sensing response. Even under the highest voltage (2 Vpp), the estimated current density at the electrode surface remains below ~1 mA $mm^{-2}$ **(Fig S6 E,F).** This low-amplitude, high-frequency regime is consistent with prior impedance-based measurements that use small alternating currents as non-invasive readouts of cell state (*18*), and with studies in excitable tissue showing that measurement-induced artifacts are strongly reduced above ~1 kHz (*19*) and effectively negligible at frequency well above 1kHz, thereby suggesting minimal perturbation of the electrophysiological state of the organoid.
